# Context-dependent presynaptic inhibition of somatostatin interneuron inputs to Layer 1 of the visual cortex

**DOI:** 10.1101/2025.09.25.678558

**Authors:** Mansour Alyahyay, Dimitri Dumontier, Deyl S. Djama, Morris Jackson, Abdullah Lugtum, Edenia Da Cunha Menezes, Haojie Ye, Jun Chen Qian, Giulia Sansone, Mohammed Soheib, Andrea M.C. Mirow, Leena Ali Ibrahim, Gabrielle Pouchelon

## Abstract

Layer 1 of the cortex is a critical site for integrating top-down inputs onto the distal dendrites of pyramidal neurons, where inhibitory neurons modulate these inputs to enable context-dependent sensory processing. Yet, it remains unclear how behavioral context dynamically regulates inhibition in Layer 1. We discover a circuit motif in which NDNF cortical interneurons (cINs) presynaptically inhibit the axonal outputs of somatostatin (SST) cINs in Layer 1 of the visual cortex. Using combinatorial genetics, monosynaptic retrograde tracing, super-resolution imaging, optogenetics, slice electrophysiology, and *in vivo* calcium imaging, we show that NDNF cINs form direct contacts onto SST axons and suppress their output, thereby modulating inhibitory responses in L2/3 pyramidal neurons. This presynaptic inhibitory circuit is preferentially engaged during locomotion and low-contrast visual conditions. By dynamically modulating Layer 1-mediated inhibitory output onto pyramidal neurons, this circuit motif enables context-dependent modulation of visual processing.

## MAIN

Layer 1 (L1) of the neocortex serves as a critical site where diverse neuronal inputs converge^1,2^. This includes bottom-up sensory inputs^3–6^, top-down inputs from higher-order cortical areas^7–10^, neuromodulatory inputs^11–13^ and cross-modal sensory inputs^14–16^. Pyramidal neurons extend extensive dendritic arborizations into L1, where they receive and integrate these inputs^17,18^, which are shaped by local cortical inhibitory neurons (cINs) to modulate their output. Within the visual cortex, arousal and locomotion strongly activate resident L1 cINs, most of which express Neuron Derived Neurotrophic Factor (NDNF) and have their axons primarily confined within L1^1,19–23^. Instead, SST cINs are activated by high-contrast visual stimuli via local recurrent connections^24–28^. The output of SST cINs, particularly the Martinotti subtype, characterized by extensive axonal arborization in L1^29–37^, targets distal dendrites of pyramidal neurons, providing recurrent inhibition, primarily via postsynaptic GABAA receptors^38,39^. In contrast, L1 NDNF cINs mediate non-specific, slow, and prolonged inhibition both via postsynaptic GABAA and GABAB receptors^40–43^. Studies using whole cell patch clamp recordings indicate that SST cINs strongly connect to NDNF cINs, while the reciprocal connection from NDNF cINs to SST somas is nearly absent^19,44,45^. However, evidence of GABA release onto presynaptic GABAB receptors at SST terminals^40,46–48^ suggests that L1 cINs could regulate SST output. However, whether NDNF cINs are specifically involved via a defined presynaptic circuit motif, and under what conditions this is engaged remains unknown.

Using in vivo calcium imaging, monosynaptic retrograde tracing, slice electrophysiology, optogenetics, and super-resolution imaging, we reveal that Layer 1 NDNF cINs form synaptic contacts onto the axons of SST cINs. This presynaptic motif contributes to the disinhibition of pyramidal neurons and is mediated in part by GABAB receptor signaling. Notably, the suppression of SST output by NDNF cINs is preferentially engaged during locomotion and low-contrast visual stimulation. Altogether, these findings identify a L1 presynaptic disinhibitory circuit motif, in which NDNF cINs regulate the output of SST cINs, providing a context-dependent basis to dynamically shape the integration of sensory inputs.

### RESULTS

#### Control of SST cIN outputs by L1 cINs

To explore whether SST cINs receive presynaptic inputs from L1 cINs, we reanalyzed our previously published monosynaptic rabies retrograde tracing dataset^49^ and quantified the number of L1 neurons retrogradely labeled from SST cINs (Fig. 1A-B). As a comparison, we also analyzed the parvalbumin (PV) cINs retrograde tracing dataset. In the visual cortex, we observed a significantly higher number of retrogradely labeled neurons in L1 from SST than from PV cINs (Fig. 1B). Retrogradely labeled L1 neurons were also detected in other cortical areas (Extended Data Fig. 1A-C). We hypothesized that these L1 retrogradely labeled cells are the main resident L1 NDNF cINs. To confirm this, we performed monosynaptic rabies tracing from SST neurons again by injecting AAV-driven helpers and rabies virus into SST-Cre mice. Using post hoc *in situ* hybridization, we found L1 retrogradely labeled neurons that expressed NDNF cIN markers, *Ndnf* and *Lamp5,* despite RNA downregulation typically induced by rabies infection^50^ (Fig. 1B, Extended Data Fig. 1D).

**Fig. 1:**
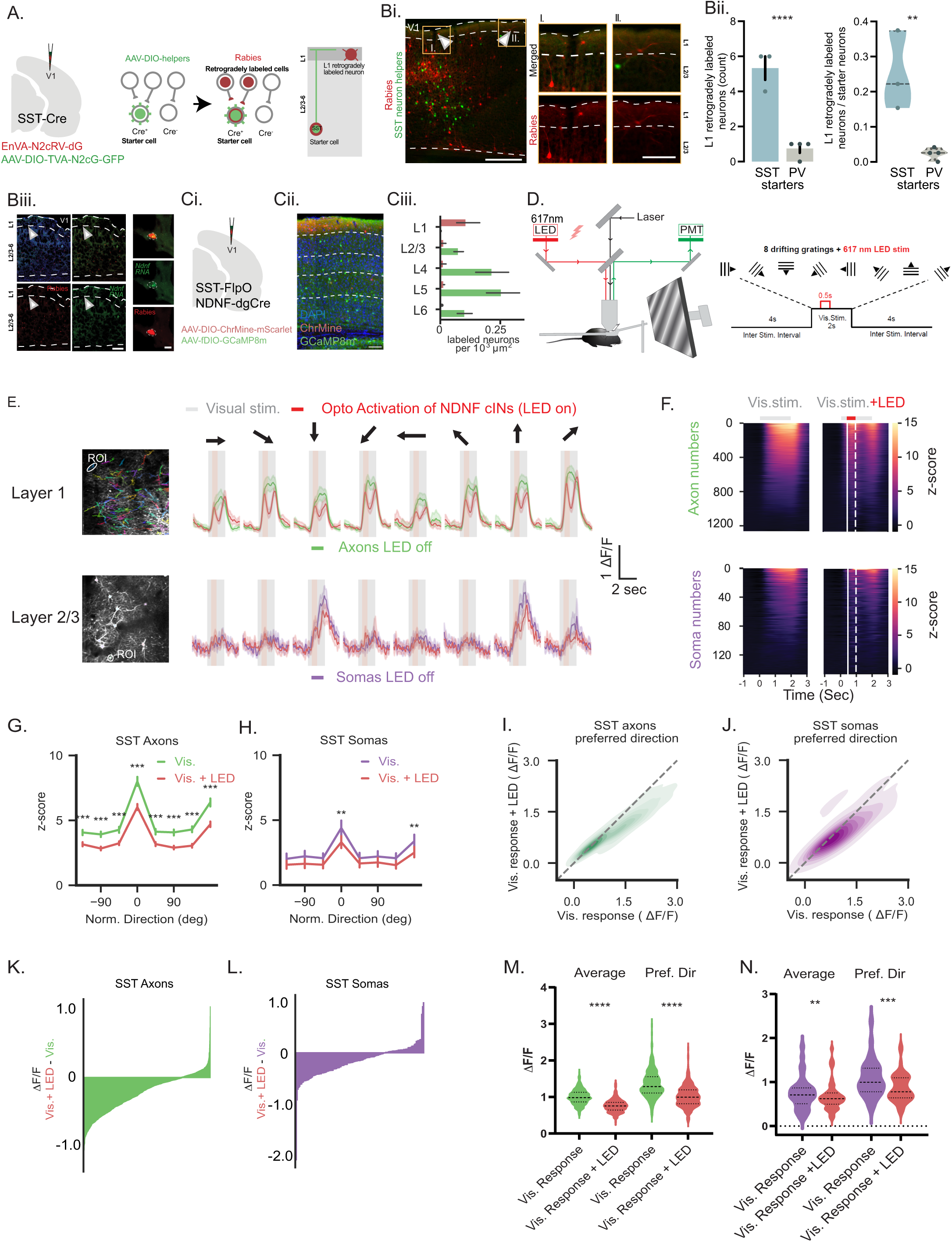
Control of SST cIN outputs by L1 cINs. **A**. Schematic of monosynaptic rabies tracing strategy from Pouchelon et al., 2021 dataset: Cre-dependent AAV-helpers containing TVA, G protein, and GFP are injected into SST-Cre transgenic mice, together with EnvA-pseudotyped CVS-N2c(ΔG) rabies virus (N2c-RV-dG, red Rabies), allowing labeling of projections to SST cINs. Quantification of retrogradely labeled cells is performed in L1 **Bi**. Example image in the visual cortex (V1) showing AAV-helper infected cells (GFP) and monosynaptically traced cells (Rabies). Arrowheads: monosynaptically traced cells in layer 1 (L1). Scale bar = 300 µm, and higher magnification of the boxed regions. Scale bar = 100 µm. **Bii.** Quantification of L1 retrogradely labeled neurons from SST and PV cINs, per mouse (N=3/3), total count (p=0.0008 t-test) and normalized per starter cells (p=0.0095, t-test). Error bars represent ±SEM. **Biii.** Monosynaptic rabies tracing from CVS-N2c (ΔG)-H2B:tdTomato and fluorescent in situ hybridization of Ndnf marker (green) in L1 (N=1 mouse). Scale bar = 100 µm, and higher magnification of the L1 neuron. Scale bar = 10 µm. **Ci**. Schematic illustration of AAV injections expressing *Cre-*dependent ChrMine and *Flp-* dependent GCaMP8m in NDNF-dgCre::SST-FlpO transgenic mice. **Cii**. Example image in visual cortex (V1) with ChRmine-expressing neurons in L1 and GCaMP8m-expressing neurons in deeper cortical layers. **Ciii**. ChrMine (red) and GCaMP8m (green) expressions across cortical layers in V1 (N = 2 mice, n = 2-3 slices). Scale bar = 200 µm. **D**. Experimental design for activating NDNF cINs and simultaneously imaging SST axons in L1 or SST somas in L2/3. **E**. Top: example ROI of an SST axon response to visual stimuli (8 directions; grey shaded area) with and without NDNF cINs activation (red shaded area). Bottom: same as top, but for SST soma. **F.** Top: responses of all ROIs of SST axons (N = 5 mice, n = 1182 ROIs) to visual stimuli (grey bar) without LED (left) and with LED (right). Solid white line: LED onset, dashed white line: LED offset. Bottom: same as top but for SST somas (N = 5 mice, n = 147 ROIs). **G**. Tuning curves of SST axon responses (z-score), aligned to each neuron’s preferred direction, with and without optogenetic activation of NDNF cINs. Green and red lines represent axon ROIs without and with NDNF cIN activation, respectively (***p<0.000125, mixed effect model). **H**. same as G but for SST somas ROIs. The purple line represents SST soma ROIs without activation of NDNF cINs (***p<0.000125, mixed effect model). Error bars represent 95% confidence intervals. **I**. Average responses of individual SST axon ROIs to visual stimuli only and to visual stimuli with the activation of NDNF cINs. **H**. Same as **I.** but for SST soma ROIs. **K**. Responses of all SST axon ROIs to optogenetic activation of NDNF cINs in all mice imaged. **L**. Same as K but for SST soma ROIs. **M**. Visual responses of SST axons in L1 were significantly lower with NDNF stimulation (Vis.Resp.: 0.81±0.034 vs. Vis.Resp.+LED: 0.66±0.029 (mean±SEM), p<0.0001, *nested t-test*). **N**. Same as K but for SST somas (Vis.Resp.: 0.82±0.13 vs. Vis.Resp.+LED:0.77±0.13, p<0.001, *nested t-test*). Error bars represent ±SEM.

A previous study has shown that monosynaptic rabies tracing can label axo-axonic circuits^51^. Given that the functional output connectivity from NDNF to SST cINs is thought to be virtually absent^24,44^, and considering the laminar segregation of their somas (L1 vs deeper layers), we hypothesized that NDNF cINs primarily inhibit SST axonal outputs in L1, a notion supported by a recently published computational model^40^. To test our hypothesis, we examined the influence of NDNF cINs on the *in vivo* responses of SST cIN output. First, we validated that NDNF cINs respond robustly to visual stimuli and optogenetic activation (Extended Data Fig. 2A-B). Next, to dissect the impact of NDNF activity on SST cIN output at different compartments, we leveraged the known anatomical organization of SST cINs. SST cINs are located in layers 2-6 with their dendrites confined near their soma. Martinotti cells, a major subtype of SST cINs, extend dense axonal arborizations into L1^31,32,52–54^. This organization allowed us to separately image SST cIN axonal activity in L1 and somatic activity in deeper layers across consecutive sessions. To achieve this, we used NDNF-dgCre::SST-FlpO mice expressing AAV-driven ChrMine^55^ and GCaMP8m^56^ in NDNF and in SST cINs respectively (Fig. 1C; Extended Data Fig. 3A-B). We recorded SST cIN responses to visual stimuli (drifting sinusoidal gratings) with and without NDNF cIN optogenetic activation in interleaved trials (Fig. 1D). Quantification from individual axonal segments and somas showed that NDNF activation significantly suppressed SST cIN responses across visual stimuli, with stronger effects in axons than in somas (Axons: Vis.Resp.: 0.81±0.034 vs. Vis.Resp.+LED: 0.66±0.029, p<0.0001; Somas: Vis.Resp.: 0.82±0.13 vs. Vis.Resp.+LED:0.77±0.13, p<0.001; Fig. 1E–J&M-N). Suppression was especially evident in the preferred direction and for the average visual response (Fig. 1M-N; Extended Data Fig. 4A–D, G–H). Additionally, a greater proportion of individual axonal segments (regions of interest, ROIs) were suppressed compared to somatic ROIs (65% axons vs 52% somas, *p*<0.005; Fig. 1K–L; Extended Data Fig. 4E–F). Altogether, these findings reveal that SST cINs receive inputs from NDNF cINs, which are engaged in an inhibitory mechanism within L1, whereby NDNF cINs suppress SST axonal output.

#### SST cINs receive presynaptic connections from L1 NDNF cINs

To elucidate whether SST axons in L1 receive anatomical inputs from L1 cINs and to further characterize the nature of this circuit motif, we analyzed a recently published serial section transmission electron microscopy (TEM) volume of mouse visual cortex acquired by the MICrONS project^57^, and leveraged the cell-type annotations generated with this dataset^58^. To identify the nature of NDNF and SST cIN connectivity, we compared these interactions with the canonical circuit formed by VIP bipolar cINs onto SST cINs. For this analysis, we selected Distally Targeting Cells (DistTC - putative SST Martinotti neurons); Sparsely Targeting Cells (SparTC - putative NDNF Neurogliaform cells) and Inhibitory Targeting Cells (InhTC - putative VIP Bipolar neurons) located only in upper layers (L1-3) (See Methods, Fig. 2A and Extended Data Fig. 5A&C).

**Figure 2:**
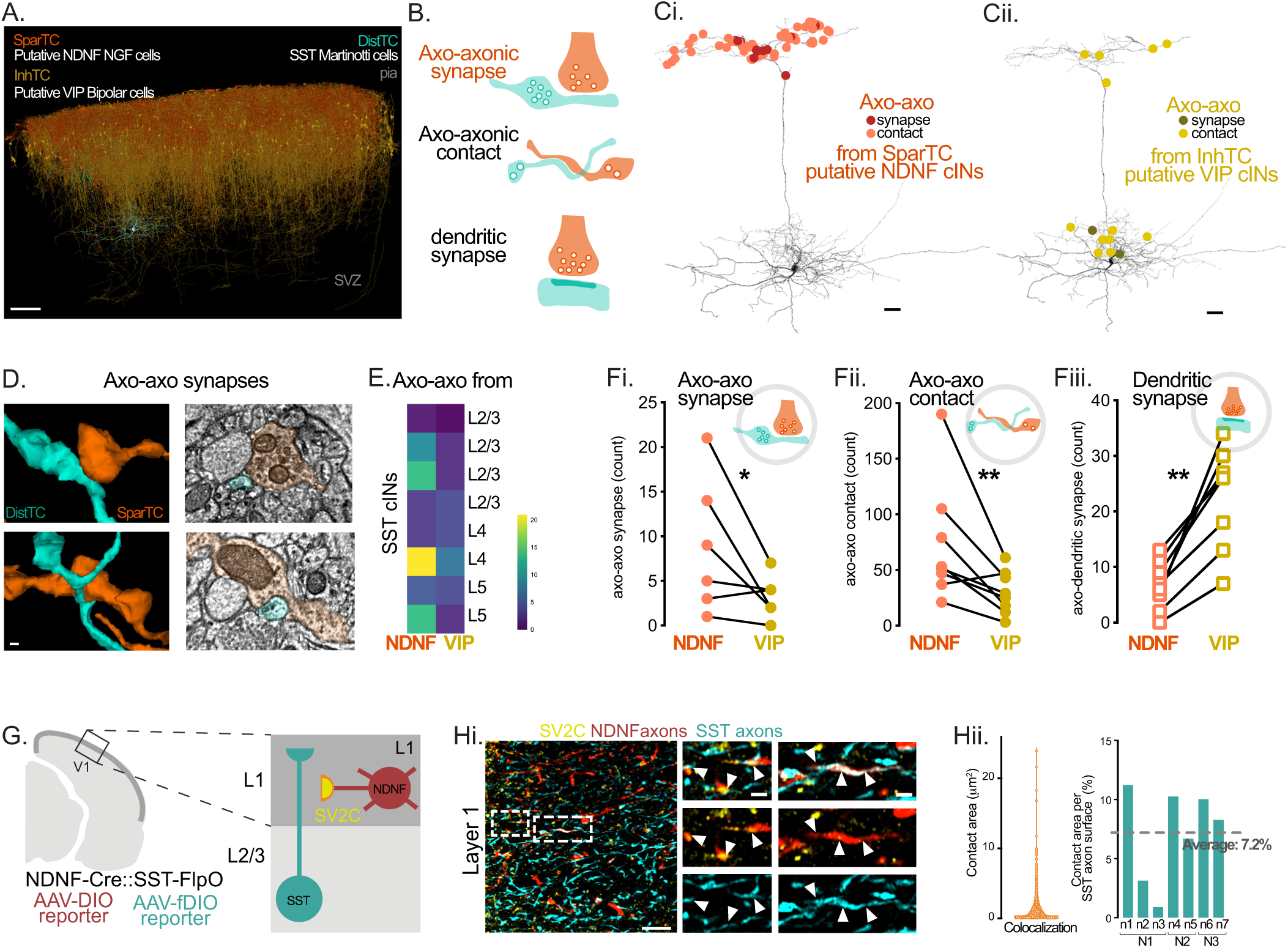
SST cINs receive presynaptic connections from L1 NDNF cINs. **A.** 3D mesh view of all upper-layer putative NDNF neurogliaform (NGC) cells, the SparTC (Sparse targeting cell), all upper-layer putative VIP bipolar cells, the InhTC, and an example of one putative SST Martinotti cell, the DistTC from the MICrONS dataset and the Schneider-Mizell et al., 2025 identification. Scale bar = 100 µm. **B.** Schematic representations of possible interactions: axo-axonic synapses, axo-axonic contacts and axo-dendritic (classic) synapses. **Ci.** Example of one DistTC (putative SST Martinotti cell), with the manually annotated axo-axonic contacts and synapses from all upper-layer SparTC. **Cii**. Example ofthe same DistTC (putative SST Martinotti cell), with the manually annotated axo-axonic contacts and synapses from all upper-layer InhTC putative VIP bipolar cells). Scale bar = 60 µm. **D.** Example of axo-axonic synapses with 3D Mesh visualization and 2D Transmission Electron Microscopy (TEM). Scale bar = 100 nm **E.** Heatmap of manually annotated axo-axonic counts per single analyzed DistTC (n = 8 neurons) organized by their soma depth (layers). **Fi.** Quantification of manually annotated axo-axonic synapses made on each DistTC from SparTC (putative NDNF cINs) and from InhTC (putative VIP cINs) (p=0.0469, One-way paired Wilcoxon). **Fii.** Same for axo-axonic contacts (p=0.0078, One-way paired Wilcoxon) **Fiii.** Same, but for automated detection from Schneider-Mizell et al. 2025 of axo-dendritic (classic) synapses (p=0.0039, One-way paired Wilcoxon).**G.** Schematic diagram of NDNF and SST cIN labeling for super-resolution imaging of axo-axonic interactions. Sections were processed from brains recorded in Fig. 2 *in vivo* imaging, with AAV-driven reporters expressed upon SST-FlpO and NDNF-dgCre. Immunostaining for SV2C was utilized to identify presynaptic terminals from NDNF axons. **Hi.** Example of staining using super-resolution AiryScan confocal imaging. Scale bar = 5 µm. Higher magnification on segments. Scale bar = 1 µm. White arrowheads: colocalization of SV2C+ NDNF and SST axons. **Hii.** Quantification of contact area surfaces, revealing large objects in addition to smaller, synaptic-like ones. Quantification of contact number between SV2C+ NDNF objects and SST axons across all replicates using super-resolution AiryScan confocal imaging and Imaris-based analysis, normalized by SST surface of axons (%) per slice and animal (N = 3 mice, n = 7 slices).

As expected for the canonical disinhibitory circuit, we found a significantly higher number of conventional axo-dendritic synapses from VIP-InhTC to SST-DistTC compared to NDNF-SparTC (Fig. 2F, Extended Data Fig. 5D-E). One limitation of the machine learning algorithm used in the MICrONS studies to annotate connections is that it detects typical features of axo-dendritic synapses, mainly thick postsynaptic density (PSD), however, atypical axo-axonic synapses are not detected^58^. Therefore, we manually identified axo-axonic interactions (See methods for details) and found a significantly higher number of contacts from NDNF cINs onto SST axons in L1, compared to the ones from VIP cINs (axo-axonic synapses from NDNF 8.75±2.48, from VIP: 3.12±0.74 per SST cell; p<0.05; Fig. 2A-F), suggesting that L1 NDNF cINs are primarily engaged in a presynaptic inhibitory circuit onto SST cIN axons. Interestingly, we identified two distinct types of axo-axonic interactions^59^: 1) axo-axonic synapse-like contacts, characterized by the presence of presynaptic vesicles, occasionally mitochondria in NDNF cINs and thin PSDs in SST cINs; 2) extended axo-axonic appositions, in which NDNF and SST axons interact over larger membrane surfaces with boutons from both cell types often located at discrete sites adjacent to the apposition zone (Fig. 2B-F; Extended Data Fig. 5C, Supplementary Videos, Supplementary Table 1).

To confirm whether these contacts are formed by genetically identified NDNF and SST cINs, we performed *post hoc* super-resolution imaging using AiryScan on brains that had been previously imaged *in vivo,* expressing fluorescent reporters in NDNF and SST cINs as described above (Fig. 1C). We identified NDNF boutons in L1 using immunostaining for synaptic vesicle protein 2C (SV2C), a presynaptic vesicle marker enriched in NDNF neurogliaform cells^1^, and observed appositions between SV2C^+^ NDNF presynaptic boutons onto SST axons. These appositions ranged from punctate to broad contact surfaces (average surface area of synaptic contacts per SST axons: 7.2%; Fig. 2G-H, Extended Data Fig. 5F), consistent with the contact types identified in the MICrONS-based analysis and supporting synaptic interactions between NDNF cINs and SST axons.

#### NDNF cINs presynaptically modulate inhibitory inputs onto pyramidal neurons

To assess whether this L1 circuit motif affects inhibition in pyramidal neurons, we tested whether the activation of NDNF cINs modulates their inhibitory inputs. As distal inhibition onto pyramidal neurons is primarily mediated by SST Martinotti cells, we recorded inhibitory postsynaptic currents (IPSCs) in L2/3 pyramidal neurons while delivering electrical stimulation ≥ 100μm away within L2/3, to primarily recruit SST-mediated distal inhibition (eIPSC) while avoiding direct activation of L1-resident cINs. Recordings were performed in the presence of excitatory synaptic blockers to isolate GABAergic transmission (Fig. 3A). We then measured evoked IPSCs by optogenetic stimulation of NDNF cINs alone (oIPSC) and together with electrical stimulation (oeIPSC). Since NDNF cINs also directly inhibit pyramidal neurons and their activation alone generates robust inhibition, we computed corrected traces for combined optogenetic/electrical stimulation (C-oeIPSCs) by subtracting the closest matching NDNF-evoked oIPSC from each trial (see Methods; Fig. 3B). We observed a significant reduction in the average IPSC peak amplitude in pyramidal neurons during NDNF cIN activation compared to electrical stimulation alone (C-oeIPSC vs. eIPSC, 17.26% average decrease; p<0.001; Fig. 3C). Notably, the magnitude of this suppression positively correlated with the amplitude of the eIPSC evoked by electrical stimulation, but not with NDNF-mediated oIPSC amplitude (Extended Data Fig. 6), likely reflecting SST-mediated inhibition. These data suggest that GABA released by NDNF cINs can tune the distal inhibition received by L2/3 pyramidal neurons.

**Figure. 3:**
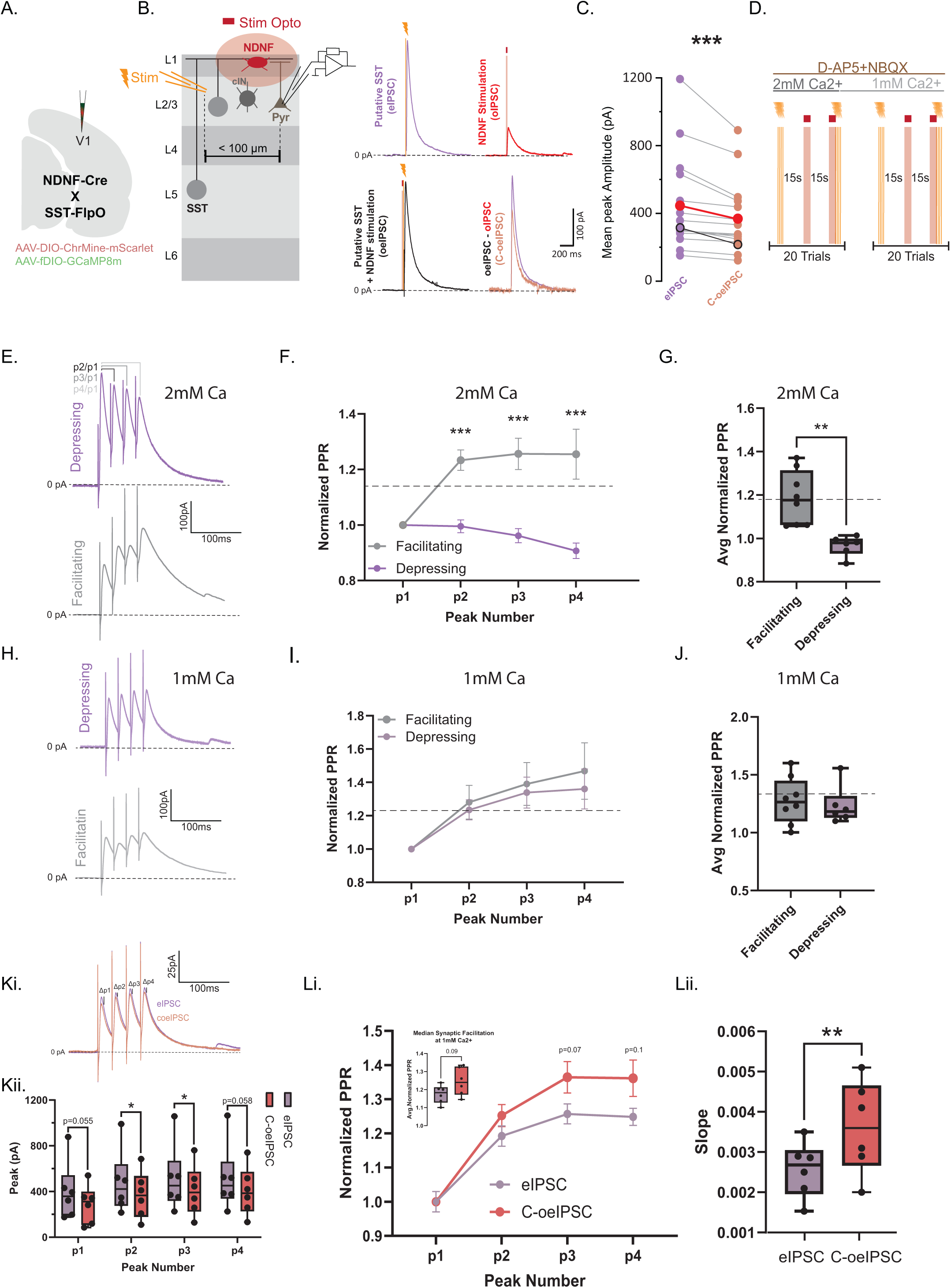
NDNF cINs presynaptically modulate inhibitory inputs onto pyramidal neurons. **A.** Schematic illustration of AAV injections expressing *Cre-*dependent ChrMine and *Flp-*dependent GCaMP8m in NDNF-dgCre::SST-FlpO transgenic mice. **B.** Left panel, illustration of the experiment where electrically and optogenetically evoked IPSCs are recorded from L2/3 pyramidal neurons. Right panel, unitary example of IPSCs evoked by minimal electrical stimulation of distant L2/3 cINs (purple trace), by optogenetic activation of local NDNF cells (red trace) and combined optogenetic followed by electrical stimulation (black trace). The NDNF component of the combined IPSC is subtracted and gives a corrected C-oeIPSC (orange trace). **C.** Plot of the mean peak amplitude for the eIPSC (in purple) and C-oeIPSC (in orange) of each recoded L2/3 pyramidal neurons showing a significant reduction in the peak amplitude when eIPSC occurred after the optogenetic activation of NDNF cells (***p=3.05e-5, n=15, Paired Wilcoxon Signed-Ranked Test for eIPSC>C-oeIPSC, 17.26% average decrease). **D.** Sequential trials of four electrical stimulation in L2/3 of 1ms pulse width are delivered at 20Hz, followed by 200ms optogenetic activation, and finally, concurrent electrical stimulation and optogenetic activation for 15-20 sweeps each at 15s intertrial interval in normal and low Ca^2+^ ASCF. >3 minute rest period is observed after changing recording conditions. **E.** Averaged current trace of pyramidal neuron recordings in response to 20Hz electrical stimulation, showing progressive synaptic depression (top) and facilitation (bottom) in Normal ACSF. **F.** Connected line graph showing the normalized paired-pulse ratio (PPR) for each peak across the eIPSC train for two population of pyramidal neurons in Normal ACSF (Depressing n=6; Facilitating n=8; Two-sample t-test (Welch Corrected); *Peak2* ***p=1.71e-4; *Peak3* ***p=8.36914e-4; *Peak4* ***p=0.00561). **G.** Box-and-whisker plot showing a significant difference in the PPR across all peaks between the two identified populations in normal ACSF (Two-sample t-test (Welch Corrected); **p=0.001). **H.** Averaged current traces of both populations of pyramidal neuron recordings in response to 20Hz electrical stimulation showing progressive synaptic facilitation in low Ca^2+^ ACSF. **I.** Connected line graph showing the normalized PPR for each peak across the electrical stimulation train for two identified populations of pyramidal neurons in low Ca^2+^ (Mann-Whitney U Test; *Peak2* p=0.47; *Peak3* p=0.47; *Peak4* p=0.65). **J.** Box-and-whisker plot showing a significant difference in the PPR across all peaks between the two identified populations in low Ca2+ ACSF (Mann-Whitney U Test; p=0.56). **Ki.** Averaged current trace of pyramidal neuron recordings in response to 20Hz electrical stimulation (eIPSC), and corrected, combined 20Hz electrical stimulation and NDNF optogenetic stimulation (C-oeIPSC; see methods). **Kii.** Box-and-whiskers plot showing the peak difference across all four peaks between eIPSC and C-oeIPSC conditions for all cells. A significant difference is found at peaks 2 and 3 (Paired Sample t-test; *Peak1* p=0.055 (12% median reduction); *Peak2 \**p=0.046(13% median reduction); *Peak3 \**p=0.046 (13% median reduction); *Peak4* p=0.058 (14% median reduction)). **Li.** Connected line graph showing the normalized PPR for each peak across the electrical stimulation train for eIPSC and C-oeIPSC conditions (Paired Sample t-test; *Peak2* p=0.137; *Peak3* p=0.07; *Peak4* p=0.11). Box-and-whiskers plot inset shows the median synaptic facilitation across all four peaks (Paired Sample t-test; p=0.09) **Lii.** Box-and-whiskers plot showing a significant increase in the rate of change of synaptic facilitation as measured by the PPR between peak1 and peak2 in C-oeIPSC conditions (Paired Sample t-test; **p=0.009). All box-and-whiskers plots show the data range, median and 1st and 3rd quartiles.

To further validate the presynaptic nature of this modulation, we examined the presynaptic probability of release using trains of electrical stimulation. A recent study demonstrated that trains of SST cIN stimulation generate depressing and facilitating responses in pyramidal neurons in the presence of standard and low calcium ACSF respectively^47^. We therefore applied trains of electrical stimulation at 20 Hz in standard (2mM Ca^2+^) and in low calcium ACSF (1mM Ca^2+^) to select for putative SST mediated distal inhibition, and we examined its modulation by NDNF cIN optogenetic stimulation (Fig. 3D). In standard ACSF, we observed two types of responses in pyramidal neurons; those that exhibit progressive depression, characteristic of SST cIN-mediated responses in pyramidal neurons; and those that exhibit facilitation as measured through their paired-pulse ratio (PPR). We found ∼43% of the neurons exhibited depressing responses (n_depressing_=6; n_facilitating_=8; p<0.001; Fig 3E-G), in line with the connection probability previously reported for L2/3 SST Martinotti cells to pyramidal neurons (Jiang et al., 2015). We used this property as a selection criterion for putative SST-mediated inhibition in pyramidal neurons. In low Ca^2+^ ACSF, the selected neurons (initially depressing) switched to a facilitating synaptic response due to lower release probability (Fig. 3H-J). Upon optogenetic activation of NDNF cINs in this condition, we observed a significant reduction in the peak amplitude of peak_2_ and peak_3_ (13% median decrease, p<0.05; Fig. 3K). Moreover, we detected an increased rate of facilitation during the initial phase of the electrical stimulation (Rate of change: eIPSC (2.57±2.8)10^-4^ vs C-oeIPSC (36±4.69)10^-4^; p<0.01; Fig. 3L), which is consistent with NDNF cINs presynaptically inhibiting the output of SST cINs on pyramidal neurons by reducing the release probability^60^. These results suggest that NDNF cINs regulate inhibition onto pyramidal neurons by presynaptically modulating SST cIN output.

#### NDNF cIN-mediated inhibition of SST output depends on GABAB signaling

Since previous studies have shown that SST Martinotti cell output is regulated by presynaptic metabotropic GABAB receptors (GABABR)^40,46–48^, we investigated whether the NDNF cIN-mediated presynaptic inhibitory motif could underlie the GABABR-dependent regulation previously described at the SST synapses. To do this, we immunolabeled GABAB1R on SV2C^+^ NDNF boutons and on SST axons defined as the postsynaptic element in the axo-axonic motif (Fig. 4A-B, Extended Data Fig. 7A). We observed that both GABAB1R-positive and -negative contacts occurred at similar densities, although GABAB1R+ contacts had significantly smaller surface areas, in line with the size of synaptic puncta (contact area GABAB1R-1.20±0.049, GABAB1R+ 0.55±0.025; p<0.0001; Fig. 4C).

**Figure. 4:**
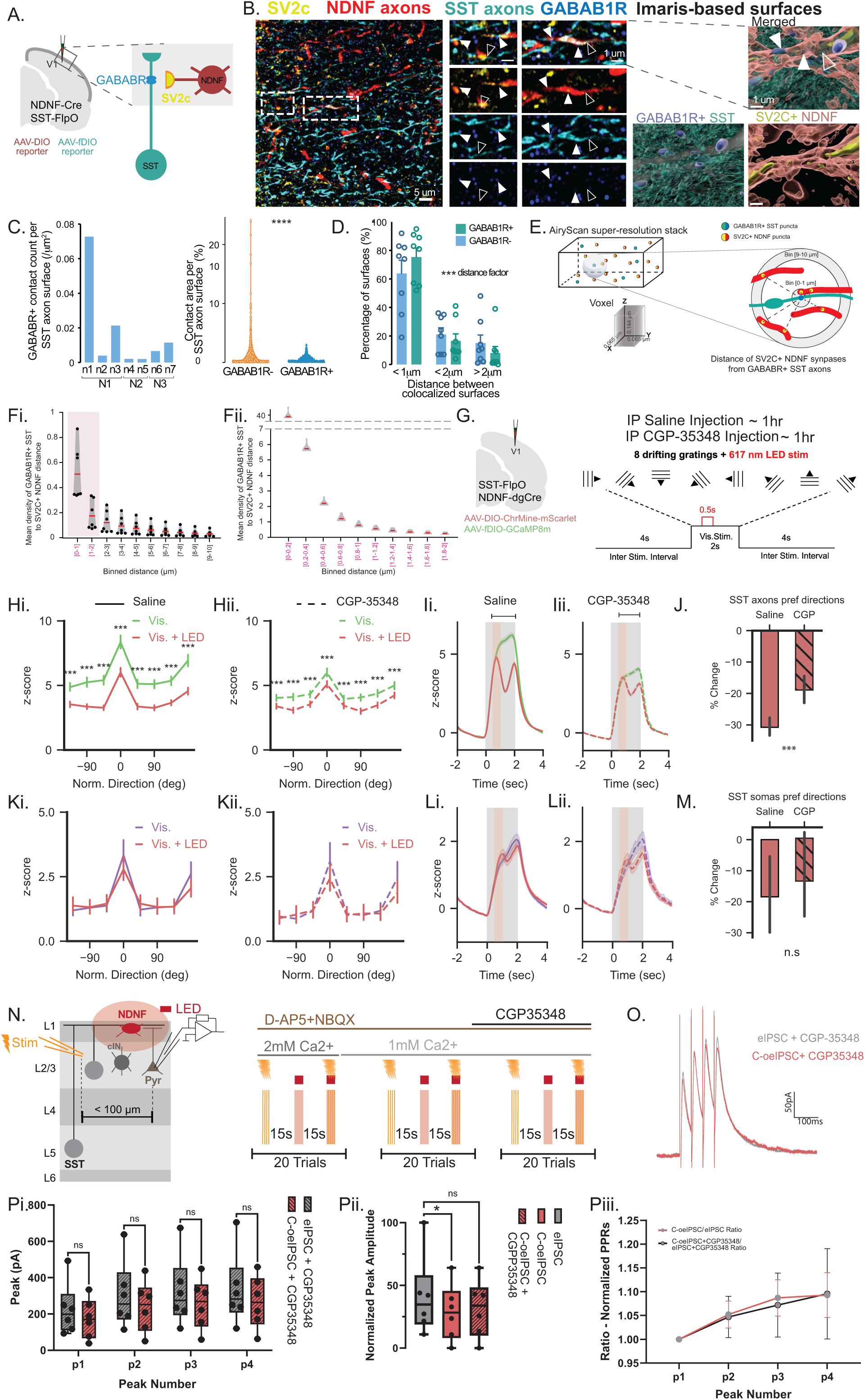
NDNF cIN-mediated inhibition of SST output depends on GABAB signaling. **A**. Schematic diagram of NDNF and SST cIN labeling in super-resolution imaging of axo-axonic interactions. Sections from Fig. 2G-H were utilized with additional GABAB1 Receptor (GABAB1R) immunostaining. **B.** Example image in layer 1 (L1) showing NDNF cIN axons (mScarlet) and SST axons (GFP) expression with labeling of presynaptic terminals (SV2c; yellow) and GABAB1R (blue) using Airy Scan, with 3D-surfaces detected using Imaris-based analysis. Scale bar = 5 µm. Higher magnification on segments. Scale bar = 1 µm. White arrowheads: colocalization of SV2C+ NDNF and GABAB1R+ SST axons. Black arrowheads: SV2C+ NDNF presynaptic terminals with close proximity to SST axons lacking GABAB1R labeling. **C.** Quantification of contact number between SV2C+ NDNF objects and SST axons across all replicates using super-resolution AiryScan confocal imaging and Imaris-based analysis, normalized by SST surface of axons (%) per slice and animal (N=3 mice, n=7 slices). Quantification of contact area surfaces, revealing large objects in GABAB1R-contacts, compared to GABAB1R+ synapses, which include more discrete puncta (n=1320/384 contacts; N = 3 mice; N = 7 slices; p<0.0001, Kolmogorov-Smirnov distribution test). **D.** Distance between GABAB1R+ SST and SV2C+ NDNF contact surfaces from the Imaris-based ‘colocalized’ population. The majority of colocalized synapses lie in <1µm distance between each other (distance factor p=0.007, GABAB+/- factor n.s., 3-way ANOVA). **E.** Unsupervised analysis of distance between SV2C+ NDNF puncta and GABAB1R+ SST puncta independently from colocalization. Distance from all SV2C+ NDNF to single GABAB1R+ SST puncta is measured and binned. **Fi.** The majority of SV2C+ NDNF are in close vicinity (0-2 µm) with GABAB1R+ SST puncta. **Fii.** Distance in higher increments from 0-2 µm bins (2x), reveals that distance mainly lies in ∼200 nm between two axons. **G**. Experimental design. After recording in saline conditions, mice were injected with the GABAB1R antagonist CGP-35348 intraperitoneally and after an hour the experiment was repeated at approximately the same field of view. In all experiments, mice were presented with moving gratings in 8 different directions (2 s; 40 trials for each direction, with 4 s inter stimulus interval) while SST INs were imaged. In half the trials, NDNF cINs were stimulated for 0.5 s using red LED. Recordings from SST axons in L1 and SST somas in L 2/3 were recorded in succession on the same day and the same field of view aside from the depth. **Hi**. Normalized responses of SST axons after saline injections (control) with visual stimulation only (green) and visual stimulation and NDNF stimulation (red) at preferred direction (n = 518 ROIs N= 5 mice; ***p<0.000125, mixed effects model). **Hii**. Normalized responses of SST axons after CGP-35348 injections with visual stimulation only (green dashed line) and visual stimulation and NDNF stimulation (red dashed line) at preferred direction (n = 426 ROIs N= 5 mice; mixed effects model). **Ii/ii**. Population response of SST axons with and without NDNF cIN optogenetic activation after treatment with saline (left) or CGP-35348 (right). Black bar: analysis window in H. **J**. Normalized difference of SST axons responses to visual stimulation in the preferred direction and NDNF stimulation in control condition (solid bar) and after CGP-35348 treatment (striped bar; saline: mean = -30.7530, 95% CI = [-33.383, -28.019], CGP: mean = -18.9620, 95% CI =[-22.933, -14.546]; ***p<0.001, mixed effects model). **K, L,** and **M**, same as H, I, and J but for SST somas (n_saline_ = 113 ROIs N_salie_ = 5 mice, n_CGP_ 87 ROIs N_CGP_ = 5 mice; saline: mean = -18.4561, 95% CI = [-31.29 -5.61], CGP: mean = -12.88, 95% CI = [-26.17 0.417]). Error bars represent 95% confidence intervals. **N.** Whole cell voltage clamp recordings are performed in L2/3 pyramidal neurons at 0mV. Sequential trials of four electrical stimulation (eIPSC) in L2/3 of 1ms pulse width are delivered at 20Hz, followed by 200ms optogenetic activation (oIPSC), and concurrent electrical stimulation and optogenetic activation (oeIPSC) for 15-20 sweeps each at 15s intertrial interval, in normal and low Ca^2+^ ASCF and in the presence of CGP35348. >3 minute rest period is observed after changing recording conditions. **O.** Averaged current traces of pyramidal neuron recordings in the presence of CGP35348 (0.1mM) responding to 20Hz electrical stimulation (eIPSC-CGP), and corrected, combined 20Hz electrical stimulation and NDNF optogenetic stimulation (C-oeIPSC-CGP). **Pi.** Box- and-whiskers plot showing the peak difference across all four peaks between eIPSC-CGP and C-oeIPSC-CGP conditions for all cells. No significant difference is found between peaks (5.5% median decrease across all peaks; Paired Sample t-test; *Peak1* p=0.077 (13% median decrease); *Peak2* p=0.08 (1% median decrease); *Peak3* p=0.11 (2% median reduction); *Peak4* p=0.16 (6% median decrease). **Pii.** Box-and-whiskers plot showing a significant reduction between the normalized peak amplitude across all four peaks for eIPSC and C-oeIPSC conditions (Paired Sample t-test; eIPSC vs C-oeIPSC p=0.049 (18% median decrease). Bath application of CGP35348 rescues the loss in peak amplitude (Paired Sample t-test; eIPSC vs C-oeIPSC-CGP p=0.14; 2% median decrease). **Piii.** Connected line graph showing the normalized PPR adjusted to the baseline condition across each peak (i.e.: C-oeIPSC/eIPSC & C-oeIPSC-CGP/eIPSC-CGP); (Paired Sample t-test; *Peak2* p=0.9; *Peak3* p=0.86; *Peak4* p=0.98).

Both contact types were primarily detected within <1 µm distance of NDNF boutons (GABAB1R+ count % at <1um: 78.71±6.13; <2um: 17.57±5.48; >2um: 3.57±1.56; p<0.001; Fig. 4D) and further unsupervised analysis revealed that the majority of NDNF boutons were positioned within 0–400 nm of GABAB1R+ SST puncta (Fig. 4E-F), consistent with the size of synaptic specializations. These findings support the presence of diverse axo-axonic interaction modes in the presynaptic modulation of SST output by NDNF cINs, including GABABR-mediated signaling.

To determine the role of GABABR signaling in the presynaptic inhibition of SST axons by NDNF cINs *in vivo*, we expressed GCaMP8m in SST cINs and Chrmine in NDNF cINs, and imaged SST axons and somas with and without NDNF activation following systemic injection of either saline or CGP, a potent GABABR antagonist^61^ (Fig. 4G). Under saline conditions, optogenetic activation of NDNF cINs suppressed SST axonal responses by 30.8% (95% CI: 28.0–33.4; *p*<0.0001; Fig. 4H-I), with 77% of axons negatively modulated (Extended Data Fig. 7B–F), consistent with the suppression observed earlier (Fig. 1E-G). Following CGP administration, suppression was significantly reduced to 19.0% (95% CI: 14.6–23.3; *p*<0.0001; Fig. 4H–J). In contrast to the axonal effects, NDNF activation did not significantly alter SST somatic responses under saline (18.5% suppression, 95% CI: 5.6– 31.3; n.s.), and CGP administration had no significant impact (12.9% suppression, 95% CI: 0.4–26.2; n.s.; Fig. 4K–M). Notably, CGP administration reduced baseline visually evoked responses in SST axons, but not in somas, consistent with a compartment-specific role for GABABR signaling in regulating axonal responsiveness (Extended Data Fig. 7C-D). Together, these results demonstrate that GABABR signaling is involved in NDNF cIN-mediated presynaptic inhibition of SST axons.

We next asked whether GABABR signaling contributes to NDNF cIN-mediated disinhibition of L2/3 pyramidal neurons. First, we confirmed the baseline involvement of GABABR in distally evoked inhibition in pyramidal neurons in acute brain slices, by recording pharmacologically isolated IPSCs evoked in L2/3 pyramidal neurons by electrical stimulation as described above (Fig. 3). baclofen, a selective GABABR agonist, nearly abolished these IPSCs (61% average decrease; baseline vs baclofen; p<0.05; Extended Data Fig. 8A-C), and this effect was reversed by addition of the GABABR antagonist, CGP (20% average decrease; baclofen vs baclofen+CGP; p<0.05). As previous studies established that baclofen does not alter SST cIN excitability^40,47^, our results suggest that GABABR regulates distal inhibitory inputs to pyramidal cells at the axonal level. With this baseline established, we next tested GABABR involvement in NDNF cIN-mediated disinhibition, using trains of electrical stimulation paired with optogenetic activation of NDNF cINs, as described above with or without CGP (Fig 4N-O). In the presence of CGP, we found that NDNF cIN activation failed to reduce the putative SST cIN inhibitory outputs to pyramidal neurons across all peaks (Fig. 4Pi; n.s). The presynaptic facilitation measured by the PPR, as observed earlier (Fig.3), was also blocked by CGP (Fig 4Piii; Extended Data Fig. 8D-E). When normalized to the first peak, we found no significant decrease in peak amplitude between electrically evoked IPSCs (eIPSC) alone and those in conjunction with NDNF optogenetic stimulation in the presence of CGP (C-oeIPSC-CGP) (Fig 4Pii; n.s). Together, these data establish that presynaptic GABABR signaling is involved in the NDNF cIN-driven suppression of SST axonal output in L1 and provide a novel mechanism for the regulation of inhibitory microcircuits in V1.

#### Presynaptic modulation of SST axons in L1 is dependent on behavioral state and visual features

We next asked under which behavioral conditions this presynaptic inhibitory circuit motif is recruited. Previous work has shown that NDNF cIN responses are modulated by behavioral states, reflected in locomotion and pupil dynamics, and that a subtype of NDNF cINs, the neurogliaform cells, are activated by low contrast visual stimuli^1,24^. Conversely, SST cINs are primarily activated by high contrast visual stimuli^24,62,63^, and while their activity is modulated by behavioral state^25,64^, this activation is weaker compared to L1 cINs^1,24^. We confirmed that indeed NDNF cINs as a general population are activated by locomotion without visual stimulation (Fig 5A-B), similar to what was previously reported for L1 cINs^1,16^ with ∼70% of the neurons exhibiting positive locomotion modulation index (LMI) (Fig 5B). Locomotion enhanced their visual responses during presentation of low contrast visual stimuli but not high contrast (Fig. 5C; Extended Data Fig. 9A-C).

**Figure 5:**
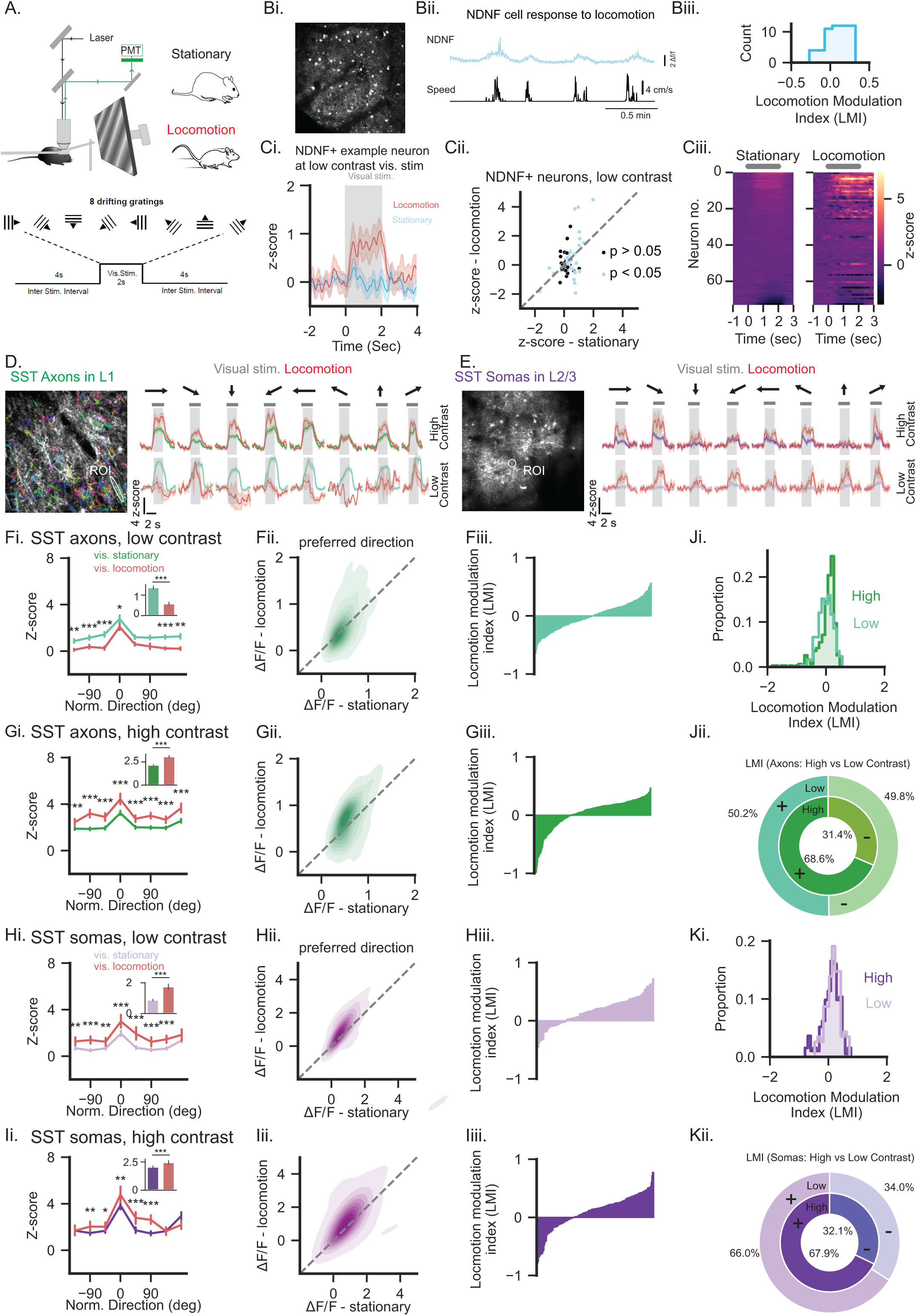
Presynaptic modulation of SST axons in L1 is dependent on behavioral state and visual features. **A.** Experimental design. Mice were habituated in the imaging setup 2 weeks after surgery where they were allowed to run freely on a linear treadmill. After habituation, sessions were recorded where mice were presented with moving gratings in 8 different directions (2 s; 10 to 40 trials for each direction, with 4 s inter stimulus interval) while SST INs were imaged. In additional sessions, mice were presented with moving gratings at low contrast (20% illumination). **Bi**. Example field of view with NDNF cINs in L1. **Bii**. Top: example NDNF cINs spontaneous activity in darkness. Bottom: corresponding locomotion speed (cm/s). **Biii.** Cumulative distribution of LMIs of NDNF cINs activity with locomotion (N=2). **Ci.** Example NDNF cIN response to low contrast visual stimuli in stationary trials (light blue) and locomotion trials (red). **Cii**. Responses of individual NDNF cINs to low contrast visual stimuli while mice were stationary and to the same stimuli with locomotion at the neuron preferred direction. **Ciii.** Normalized responses to low contrast visual stimuli of all ROIs of NDNF cINs when mice were stationary (left) and with locomotion (right; N = 2 mice, n = 73 ROIs). **D**. Left: Example field of views with SST axons in L1. Top traces: example roi of SST axon presented with gratings at high contrast. Bottom traces: example roi of SST axon presented with gratings at low contrast. Red line: responses to visual gratings during locomotion. **E.** same as D but for SST somas in L2/3. **Fi.** Normalized responses of SST axons with low contrast visual stimulation only (light green) and visual stimulation with locomotion (red) across all directions presented. Inset: Average responses across directions (n_stationary_ = 342 N = 5 n_locomotion_ = 330 ROIs N = 5; stationary: mean 1.38, 95% CI = [1.31, 1.46], locomotion: mean 0.581, 95% CI = [0.51 0.65]). **Fii.** Responses of individual SST axon ROIs to low contrast visual stimuli while mice were stationary and to low contrast visual stimuli with locomotion at SST preferred direction. The data are fit with a gaussian kernel for visualization. **Fiii.** LMI of SST axons during presentation of low visual contrast. **G, H, and I.** same as **F** but for SST axon responses at high contrast (n_stationary_ = 468 N = 6 n_locomotion_ = 436 ROIs N = 5; stationary: mean 2.18, 95% CI = [2.13, 2.23], locomotion: mean 3.09, 95% CI = [3.00, 3.20]), SST somas at low contrast (n_stationary_ = 198 N = 9 n_locomotion_ = 162 ROIs N = 8; stationary: mean 0.88, 95% CI = [0.82, 0.94], locomotion: mean 1.75, 95% CI = [1.58, 1.91]), and SST somas at high contrast (n_stationary_ = 297 N = 12 n_locomotion_ = 257 ROIs N = 11; stationary: mean 2.05, 95% CI = [1.96, 2.15], locomotion: mean 2.46, 95% CI = [2.29, 2.62]), respectively. **Ji**. Histogram of SST axons LMI with the presentation of low contrast (light green) and high contrast (dark green). **Jii.** Proportion of SST axons LMI. Outer ring (low contrast): positive LMI (green), negative LMI (light green). Inner ring (high contrast): same categories and colors as above. **K.** Same as **J**, but for SST somas.

Based on this, we reasoned that locomotion and low contrast conditions that strongly recruit NDNF cINs should result in suppression of SST axonal activity. To test this, we performed two-photon imaging of GCaMP8m-labeled SST axons in L1 and SST somas in L2/3 (Fig. 5D-E) during both low (20%) and high contrast (∼100%) visual stimulation, while monitoring the locomotion to classify responses according to running or stationary states. Under high-contrast conditions, SST axons in L1 exhibited clear orientation-tuned responses during the stationary period, which were further enhanced during locomotion (stationary: 2.18, 95% CI 2.13–2.23; locomotion: 3.09, 95% CI 3.00–3.20; p<0.0001; Fig. 5D, upper right; Extended Data Fig. 10A–C). Their LMIs were predominantly positive (68.6%; Fig. 5G-J). In contrast, at low contrast, SST axonal responses were suppressed during locomotion (stationary: 1.38, 95% CI 1.31–1.46; locomotion: 0.58, 95% CI 0.51–0.65; p<0.0001; Fig. 5D, lower left, 5F), with ∼50% of axonal ROIs showing negative LMIs at the preferred orientation (Fig. 5J). By comparison, SST somas in L2/3 consistently increased their responses with locomotion at both low and high contrasts (low: stationary 1.93, 95% CI 1.83–2.02; locomotion 2.50, 95% CI 2.32–2.69; p<0.0001; high; stationary 2.05, 95% CI 1.96–2.15; locomotion 2.46, 95% CI 2.29–2.62; p<0.0001; Fig. 5E&H-I) in line with previous studies^24,62,63,65^. Their LMIs were predominantly positive under both low (66.0%) and high (67.9%) contrast stimulation (Fig. 5K). Thus, presynaptic inhibition of SST axons in L1 is selectively engaged under the very conditions that robustly recruit NDNF cINs, namely locomotion and low-contrast visual input, and is largely bypassed when sensory drive is strong.

### DISCUSSION

Our study identifies a previously unrecognized presynaptic disinhibitory circuit motif in layer 1 (L1) of the visual cortex, in which NDNF cortical interneurons (cINs) selectively suppress the axonal output of somatostatin (SST) cINs. We demonstrate that NDNF cINs form axo–axonic contacts and GABAB receptor-positive synapses onto SST axons. Moreover, we show that NDNF cINs inhibit SST axonal output, thereby attenuating distal inhibition received by L2/3 pyramidal neurons. This suppression occurs during locomotion and low contrast visual stimulation, conditions that robustly engage NDNF cINs. These findings establish a novel presynaptic inhibition mechanism that can adjust the strength of SST cIN-mediated inhibition, enabling flexible integration of top-down and bottom-up signals and fine-tuning visual responsiveness.

Inhibitory control of SST cINs has primarily been described in the motif involving VIP cINs, in which cholinergic or locomotion-dependent recruitment of VIP cINs suppresses SST somatic output, producing widespread disinhibition of pyramidal neurons, modulating pyramidal neuron excitability globally across the dendritic arbor^25,26,66^. VIP cINs are primarily activated under low contrast conditions, but they lack orientation tuning and are preferentially engaged by larger visual stimuli^67^. In contrast, NDNF cINs, while also recruited at low contrast and during locomotion, exhibit orientation tuning. Given that the NDNF cIN– specific motif we describe acts presynaptically at SST axon terminals in L1, and that SST cINs can gate and compartmentalize dendritic activity in L2/3 pyramidal neurons^37,68^, this circuit could translate state-dependent NDNF cIN activity into local, orientation-selective inhibition at apical dendrites, which may sharpen orientation tuning at low contrast during locomotion.

Apical dendrites of pyramidal neurons are major sites of top-down and modulatory input integration, whereas bottom-up sensory drive primarily targets basal dendrites and perisomatic regions. Driven largely by recurrent excitation from bottom-up sensory inputs, SST cINs strongly regulate apical dendritic activity and gate the impact of top-down inputs^69^. By reducing SST output, NDNF cIN-mediated presynaptic inhibition could selectively enhance top-down influence on pyramidal neurons. This compartment-specific disinhibition could improve the signal-to-noise ratio for weak inputs, including low contrast stimuli, during locomotion, while preserving responsiveness for strong visual stimulation. Such selectivity preserves stimulus tuning while flexibly adjusting network excitability according to behavioral context, an advantage over global disinhibitory mechanisms that might also degrade stimulus selectivity. Importantly, previous studies have shown that SST cINs are the principal source of inhibition onto NDNF cINs^19,44^. By showing that the recruitment of the presynaptic inhibition motif is both state- and stimulus-dependent, our findings suggest there might be a competitive relationship in which the relative activation of NDNF and SST cINs determines the inhibitory tone at apical dendrites. Together with our data, this supports the existence of a reciprocal inhibitory motif in L1, pointing to a bidirectional interplay that dynamically regulates inhibitory signaling and tunes the balance of top-down and bottom-up integration.

Mechanistically, presynaptic inhibition has been shown to involve both GABAA and GABAB receptors^40,70–72^. Here, we show that this presynaptic disinhibition involves GABAB receptor signaling. We observed GABAB1R-positive puncta at NDNF–SST axo–axonic contacts, and pharmacological blockade with CGP attenuated NDNF cIN-evoked suppression of SST axons, but not in somas. However, 18% of the NDNF cIN mediated suppression was not blocked by CGP, suggesting the presence of other GABABR negative synapses, potentially driven by GABAAR. Consistent with this diversity of synapses, MICrONs data analysis revealed distinct modes of synaptic contacts. Furthermore, CGP-blockade was the strongest during the NDNF cIN activation (during the 0.5s of LED stimulation), while weaker after the cessation of the LED stimulation, suggesting that both fast and slow modes of neurotransmission exist. In addition, we observed that CGP administration reduced baseline visually evoked responses selectively in SST axons, but not in somas, consistent with a compartment-specific contribution of GABABR signaling to axonal responsiveness. This raises the possibility that, beyond NDNF cINs, additional GABAergic inputs or neuromodulatory mechanisms might engage axonal GABABRs to dynamically regulate SST output. It is possible that neurogliaform cells in other layers could be engaged in this circuit. Identifying these potential convergent sources of presynaptic inhibition will be important for determining how L1 circuits integrate multiple inhibitory influences and coordinate gain control across behavioral states.

Moreover, the coexistence of GABABR-positive and –negative contacts and structurally diverse NDNF➔SST inhibitory synapses, raises the possibility of parallel mechanisms of presynaptic modulation. Indeed, only part of the SST cIN population is suppressed in these conditions, suggesting specific organization within SST cIN subtypes. SST cIN subpopulations are defined by their local input/output relationship and their laminar positions within the cortex^36^. The electron microscopy data analysis suggests a potential gradual increase of axo-axonic contacts with deep-layer SST cINs, supporting this hypothesis. Layer-specific inhibitory motifs offer parallel routes for controlling pyramidal neuron output. VIP cINs act postsynaptically on distinct subtypes of SST cINs, providing global disinhibition, while NDNF cINs act presynaptically on SST axons, offering localized, input- and layer-specific disinhibition. However, it is yet unclear whether these two disinhibitory motifs interact with distinct SST cIN subtypes in different layers, and whether distinct SST cIN subtypes are engaged in specific context-dependent modulations. Addressing this would demonstrate how the coexistence of both disinhibitory strategies expands the computational power of cortical circuits, enabling modulation across behavioral, spatial and temporal scales. This arrangement could support circuits to dynamically reconfigure according to sensory reliability and behavioral demands.

By uncovering a presynaptic disinhibitory motif, our findings provide a new mechanism for top-down and bottom-up signal integration. This circuit enables context-dependent reweighting of inputs at a subcellular level, offering a means to prioritize behaviorally relevant information without altering network excitability globally. Future work should explore the context driving similar presynaptic motifs in other sensory cortices, including the somatosensory cortex, and higher-order areas, such as the anterolateral medial (ALM) cortex, as we also revealed retrogradely labeled L1 cIN from SST cIN in those areas. Since neuromodulatory systems are engaged in disinhibition, it would be of interest to examine whether they modulate the NDNF cIN-mediated presynaptic inhibition of SST axons. Determining how this microcircuit is altered in disease models with disrupted GABABR signaling or impaired L1 cIN function could reveal novel therapeutic targets for disorders of sensory integration.

## ACKNOWLEDGEMENTS

We thank Sam Liebman (CSHL) and Nor Linda Abdullah (KAUST) for help with genotyping and maintaining the animal colony. We thank Dr. Qing Xu and Mashael Felemban for custom AAV viral production at KAUST. We thank the director of the CSHL Microscopy core Facility, Dr. Erika Wee, for the access to the LSM980 AiryScan confocal, and advice on high-resolution imaging. We thank Prof. Christian Froekjaer Jensen, Prof. Jesper Tegner, Dr. Jennifer Sun, Dr. Thomas Younts and members of the Ibrahim and Pouchelon labs for their valuable comments on the manuscript. This work was supported by Pershing Square Innovation Fund (PSIF) to GP and Baseline Research Fund and KAUST Competitive Research Grant 11 to LAI. The CSHL microscopy core facility is funded by: S10 grant number (NIH S10OD034372).

## CONTRIBUTION

Conceptualization: LAI and GP; Experiments: In vivo imaging & visual stimulation: MA, MS. Surgeries: MJ, AL, MA. Patch clamp recordings: DSD, DD. Histology and Imaging: HY, MJ and AL. Formal Analysis: MA, DSD, DD, GS, EDM, JQ, HY, LAI, GP. Writing and editing: all authors. ACMS was involved in the early phase of the project.

## LEGENDS

**Extended Data Fig. 1:**
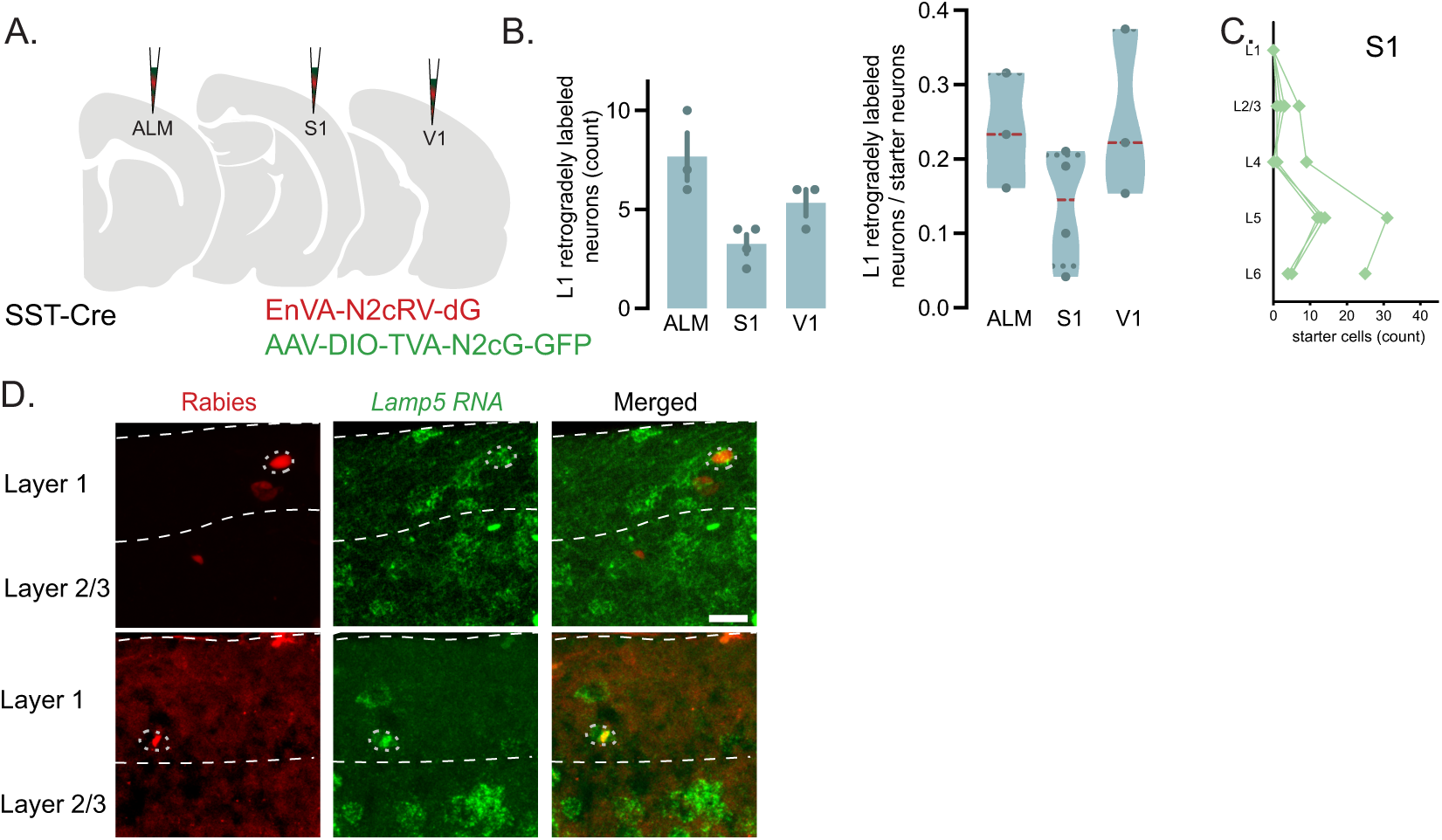
Monosynaptic retrograde rabies tracing of L1 neurons from SST cINs. **A**. Areal specific schematic of monosynaptic rabies tracing strategy from Pouchelon et al., 2021 dataset: Cre-dependent AAV-helpers containing TVA, G protein, and GFP are injected in Anterolateral Motor Cortex (ALM), the primary Somatosensory (S1) and Visual (V1) cortices of SST-Cre transgenic mice, together with EnvA-pseudotyped CVS-N2c(ΔG) rabies virus (N2c-RV-dG, red Rabies), allowing labeling of projections to SST cINs. **B.** Quantification of L1 retrogradely labeled neurons from SST in ALM, S1 and V1, per mouse (N=3/3), total count and normalized per starter cells. Error bars represent ±SEM. **C.** Distribution of starter SST cIN in V1 across layers. **D.** Monosynaptic rabies tracing from CVS-N2c(ΔG)-H2B:tdTomato and fluorescent in situ hybridization of *Lamp5* marker (green) in L1 (N=2 mice). Scale bar = 20 µm.

**Extended Data Fig. 2:**
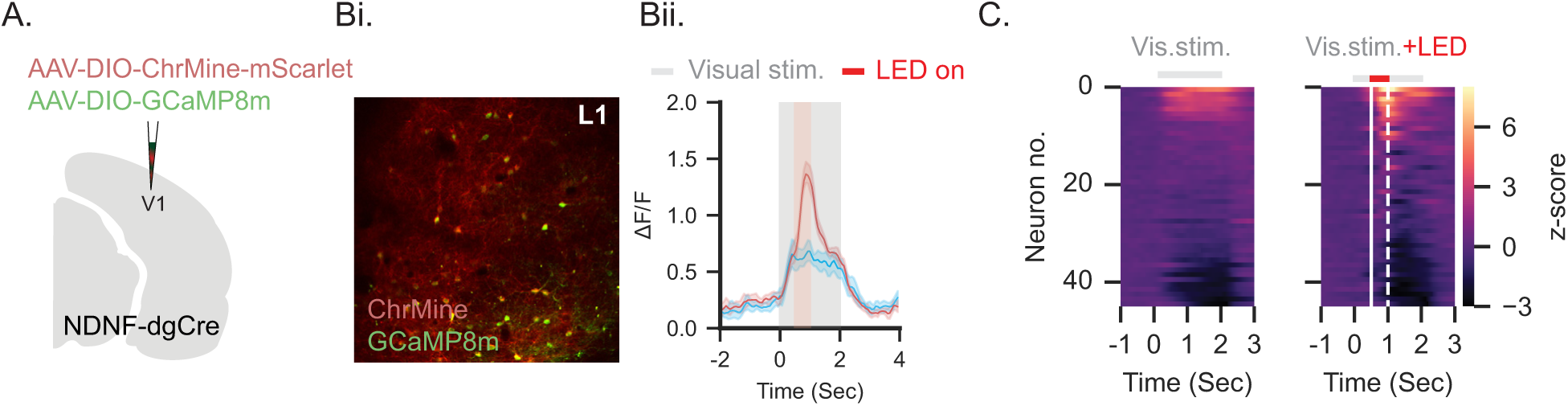
Responses of NDNF cINs to optogenetic activations. **A**. Schematic illustration of AAV injections expressing *Cre-*dependent ChrMine and *Cre-* dependent GCaMP8m in NDNF-dgCre transgenic mice. **B**. **i**. Example field of view of NDNF cINs expressing GCaMP8m and ChrMine in L1 (N = 2 mice). **ii**. Example of NDNF cINs with and without optogenetic activation. C. All ROIs of NDNF cINs (n = 45 ROIs) without LED (left) and with LED (right). Grey bar: visual stimuli, solid white line: LED onset, dashed white line: LED offset.

**Extended Data Fig. 3:**
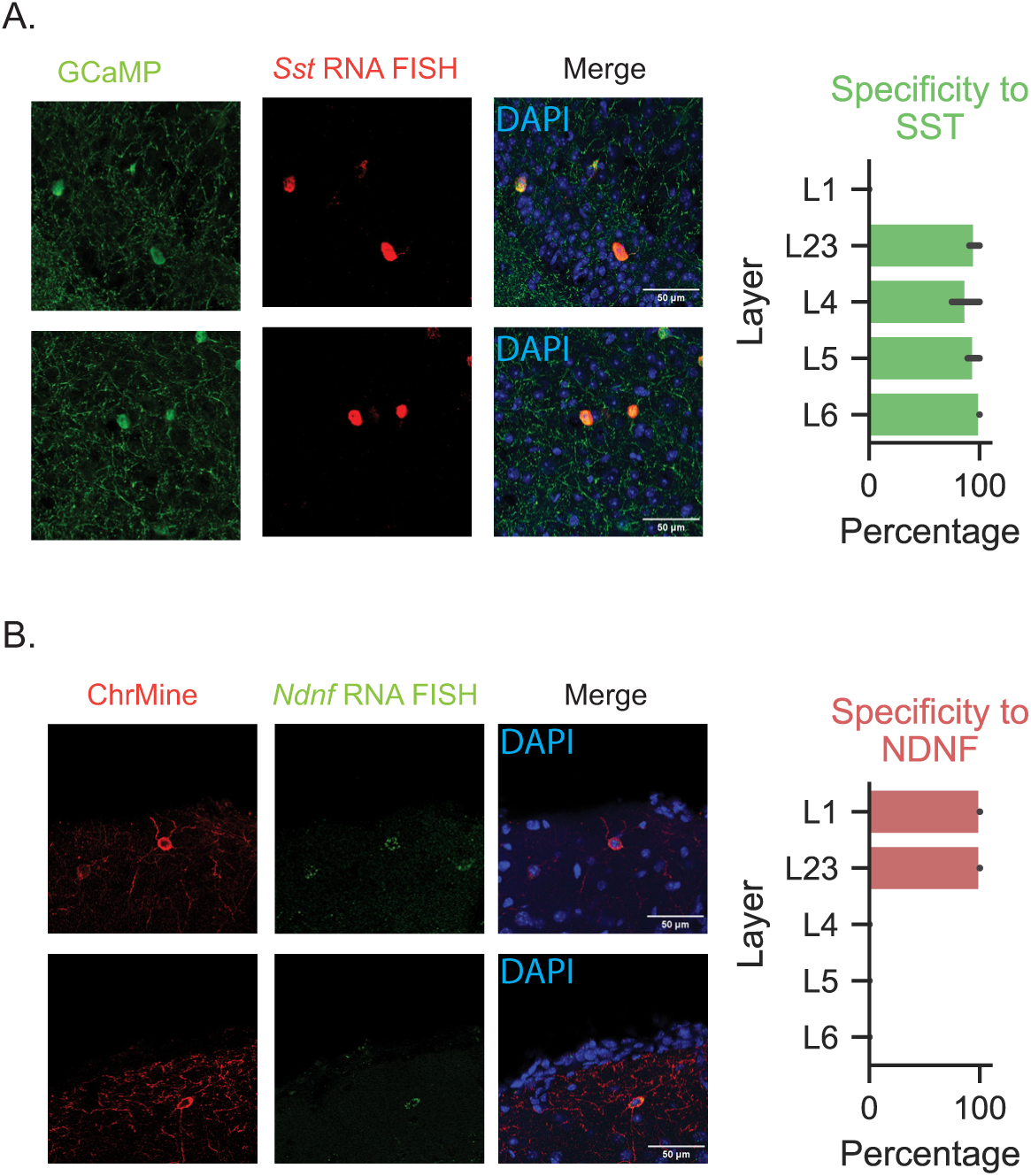
Specificity of GCaMP8m and ChrMine expression. **A**. Left: Coronal sections of GCaMP8m expression (green) and *Sst* RNA ISH labeling (red). Right: percentages of GCaMP8m specificity to SST cINs across layers in the visual cortex (N = 2 mice). **B**. Same as A, but for ChrMine expression and NDNF cINs. Error bars represent 95% confidence intervals.

**Extended Data Fig. 4:**
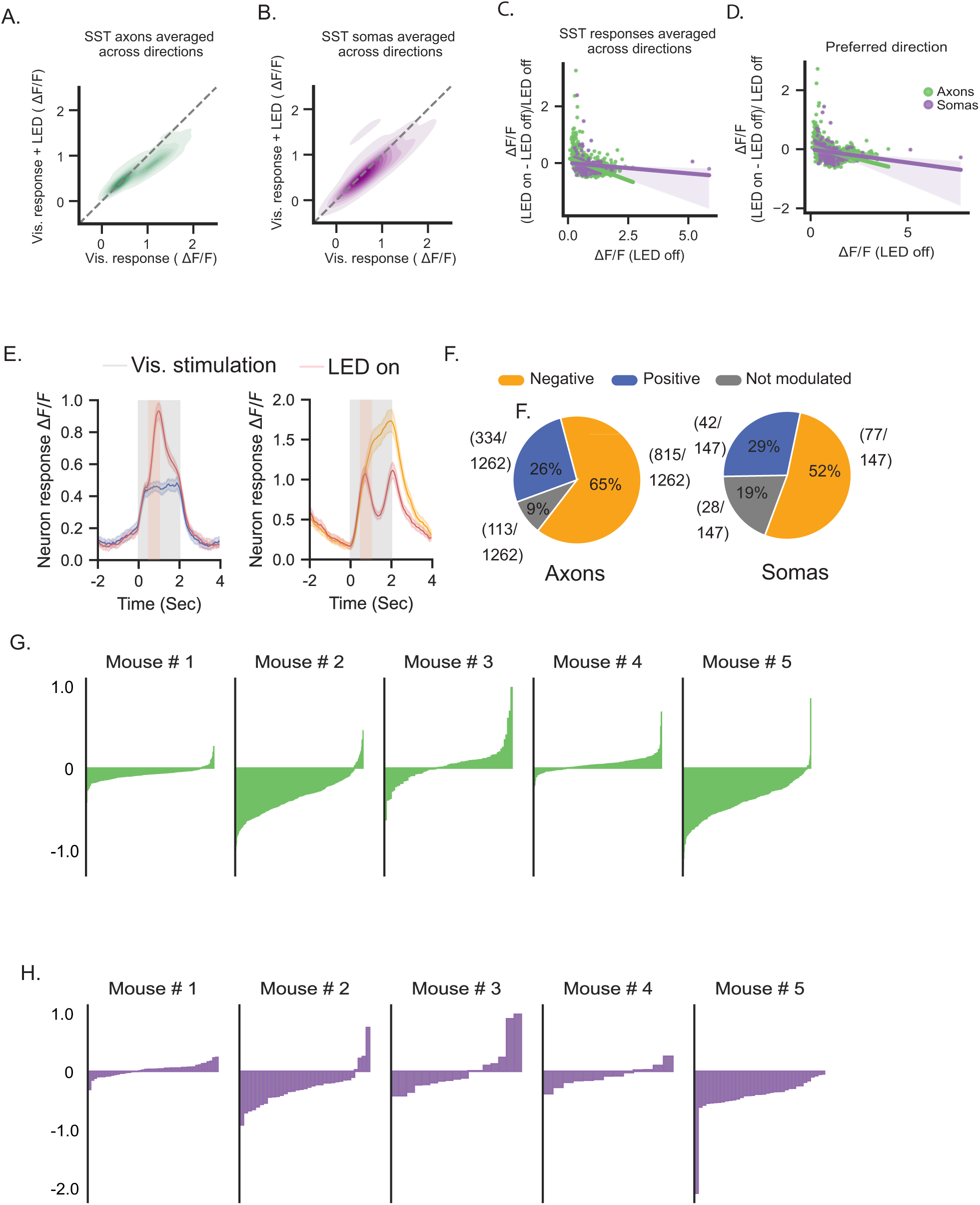
Effect of NDNF cINs activation on individual SST axons and somas. **A**. Responses of individual SST axon ROIs to visual stimuli only and to visual stimuli with optogenetic activation of NDNF cINs averaged across all directions. **B**. Same as A, but for SST somas ROIs. **C**. Linear regression model for both SST axon and soma average responses to visual stimuli, comparing conditions with and without optogenetic activation of NDNF cINs. **D**. Same as C, but in the preferred direction of each ROI. **E**. Left: example of SST axon enhanced response to visual stimulation with and without NDNF cIN activation. Right: example of SST axon suppressed by NDNF cINs activation. **F**. Left: ratios of different responses of SST axons to NDNF cIN activation. Right: same as left but for SST somas. **G**. Responses of individual mice SST axon ROIs to optogenetic activation of NDNF cINs. **H**. Same as G but for SST somas.

**Extended Data Fig. 5:**
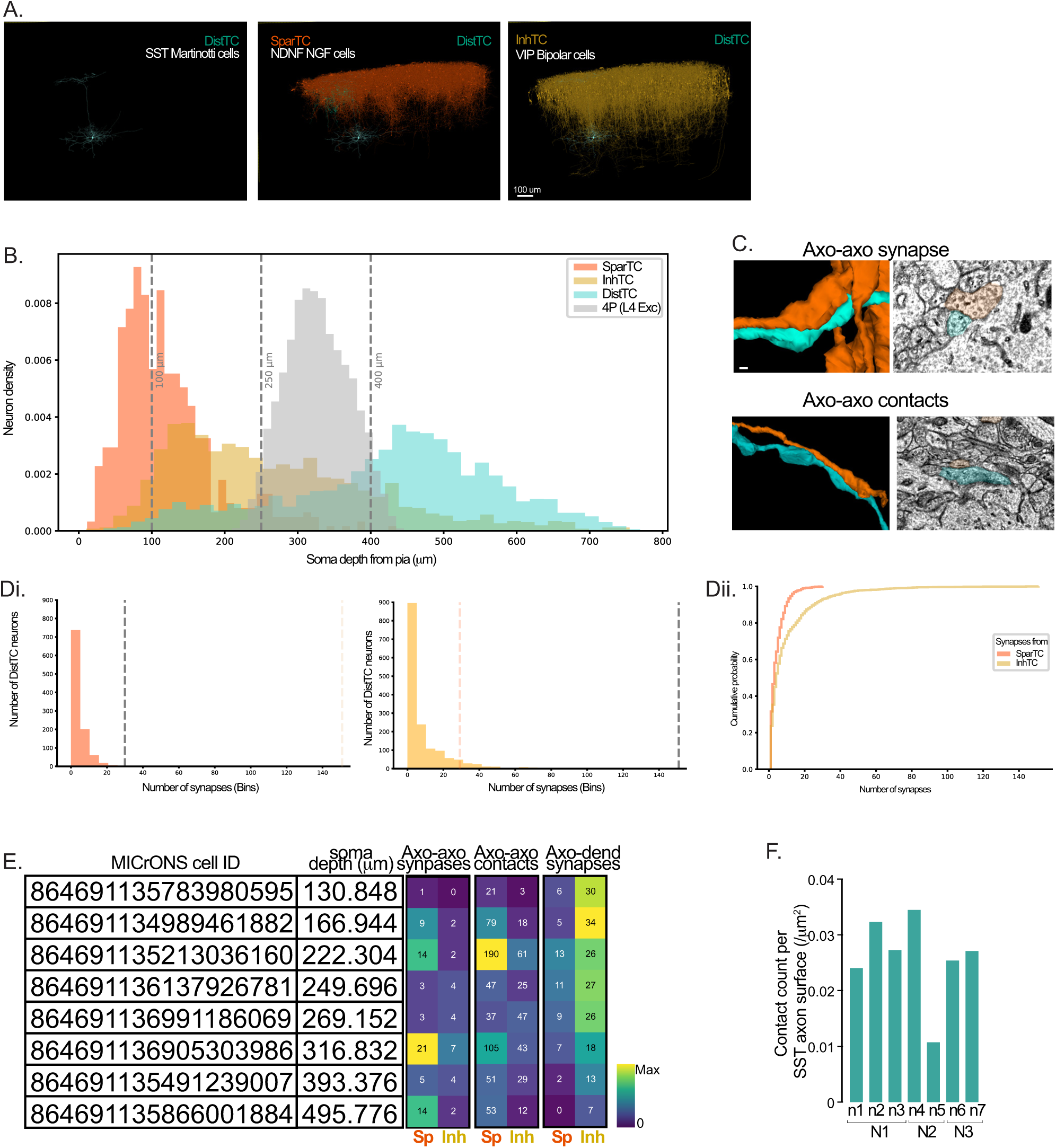
Super-resolution imaging of SST and NDNF axons. **A. A.** 3D mesh view of an example of one putative SST Martinotti cell, the DistTC, all upper-layer putative NDNF neurogliaform (NGC) cells, the SparTC (Sparse targeting cell), all upper-layer putative VIP bipolar cells, the InhTC, from the MICrONS dataset and the Schneider-Mizell et al., 2025 identification. **C.** Soma distribution per depth supporting the selection of upper-layers SparTC and InhTC, as well as identification of DistTC laminar location. All “SST” DistTC (green), “NDNF” SparTC (red), “VIP” InhTC (yellow), and “Layer 4 excitatory” 4P neurons serve as a laminar marker. Dotted grey lines highlight putative divisions of L1, L2/3, L4 and L5/6. The 250µm limit was picked to select upper-layer SparTC and InhTC. **B.** Examples of axo-axonic synapse and axo-axonic contact with 3D Mesh visualization and 2D Transmission Electron Microscopy (TEM).Scale bar = 100nm. **Di.** Number of automatically detected axo-dendritic synapses (classic synapses) formed by “NDNF” SparTC and “VIP” InhTC per DistTC. Grey dotted lines represent the maximum synapses for each condition. Colored dotted lines represent the maximum of the other conditions, for comparison. **Dii.** Cumulative distribution of the number of synapses. **E.** MICrONS-based DistTC identity of cells included in this analysis, together with their soma-depth and heatmap of counts for axo-axonic contacts, axo-axonic synapses, and axo-dendritic synapse formed by SparTC and InhTC onto DistTC. **F.** Quantification of the number of contacts between SV2C+ NDNF objects and SST axons across all replicates (N = 3 mice, n = 7 slices).

**Extended Data Fig. 6:**
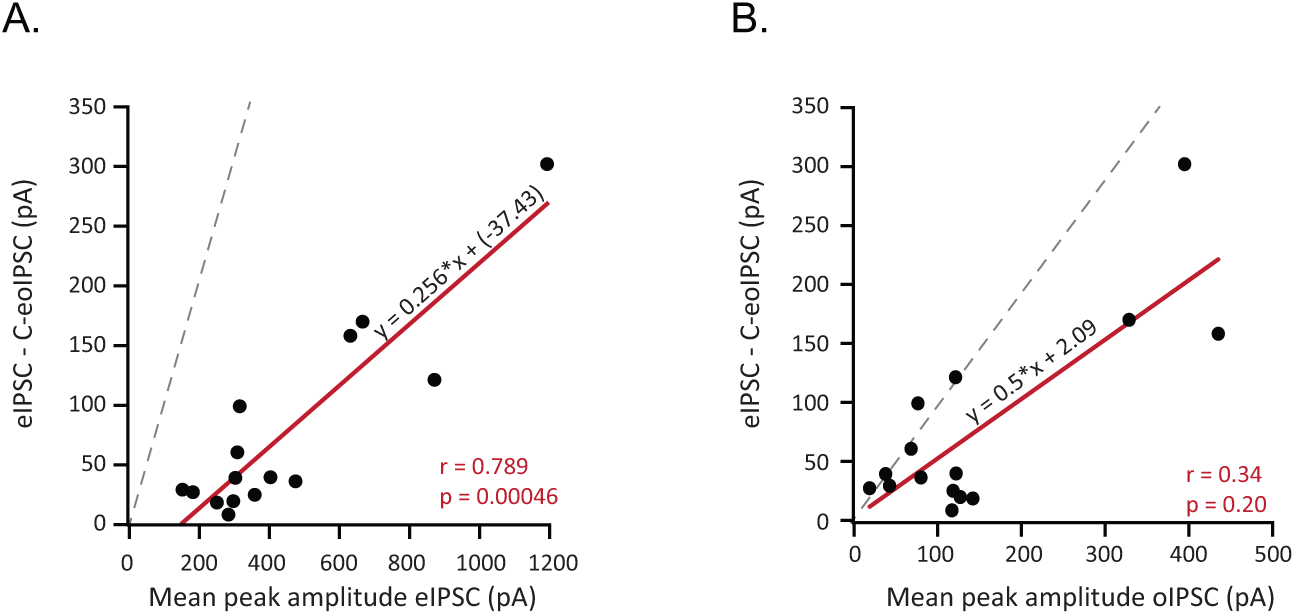
Presynaptic effect of modulated inhibition in pyramidal cells. **A.** Correlation between eIPSC - C-oeIPSC mean peak amplitudes and eIPSC mean peak amplitudes of the same L2/3 pyramidal cells (slope: 0.256, intercept: -37.43; Spearman correlation r: 0.789, p: 0.00046). **B.** Same as A between eIPSC - C-oeIPSC mean peak amplitudes and oIPSC mean peak amplitudes (slope: 0.50, intercept: 2.09; Spearman correlation r: 0.34, p: 0.20). The red lines are the linear regressions and the grey dotted lines are the diagonals.

**Extended Data Fig. 7:**
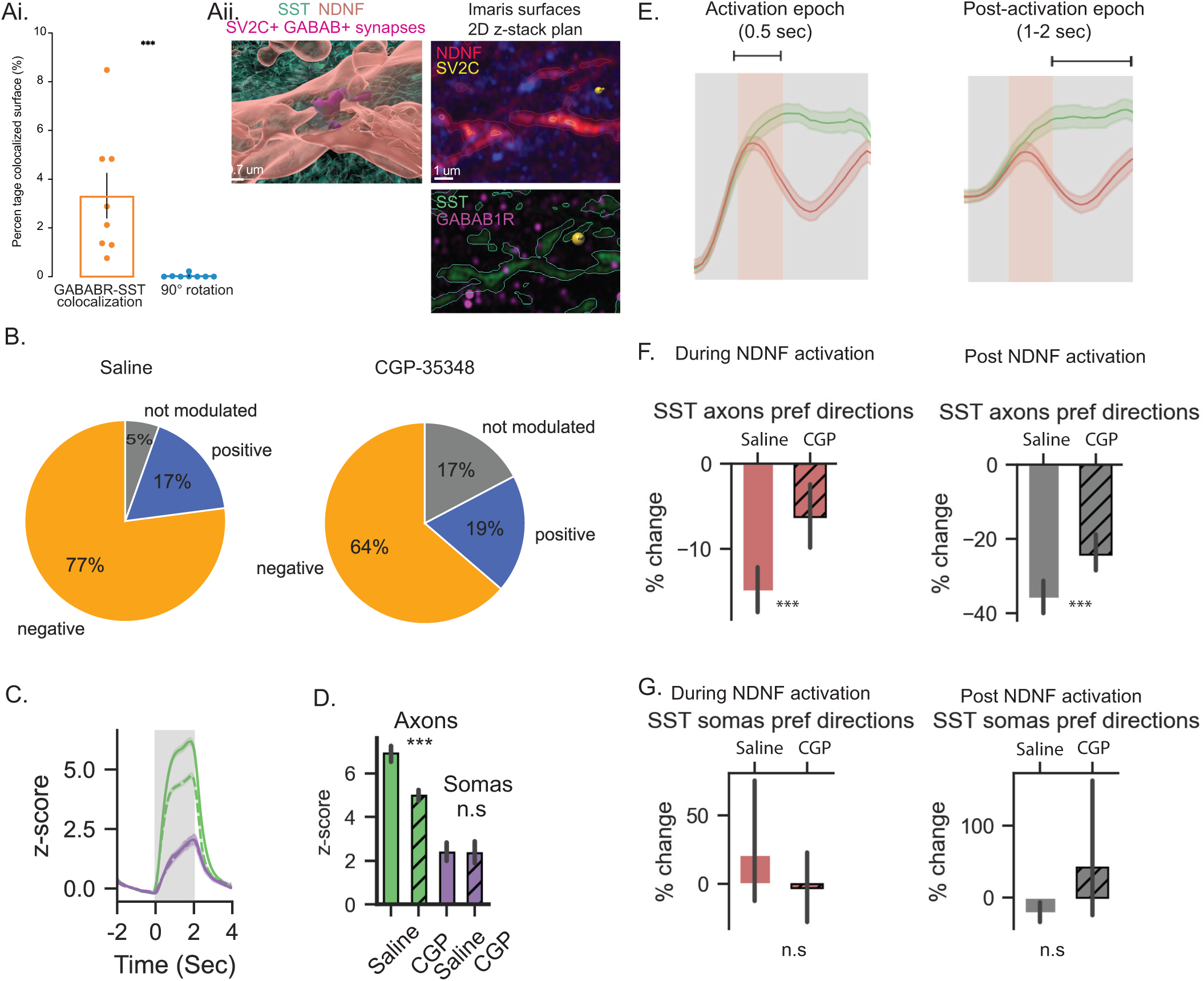
GABAB antagonist abolishes NDNF cIN suppression of SST cIN axons but not somas. **Ai**. Specificity of Imaris-based colocalization in detecting synapses. 90° rotation of GABABR1 channel significantly decreases colocalization detections with SST axons. (p = 0.0002, Mann-Whitney test). **Aii.** Left: Imaris-based surface detection (magnification of segment from Fig. 4B). SST axons (green) colocalize with GABAB1R and NDNF axons (orange) colocalize with SV2C. Colocalization of the two reveal GABAB1R+ synaptic contact (purple). Right: 2D-plan from Z-stack imaged. Surfaces are identified using Imaris. **B.** Ratios of axon ROIs modulated by the optogenetic activation of NDNF cINs with saline treatment (left) and CGP-35348 treatment (right). **C**. Population responses of SST axons and somas with saline treatment and CGP-35348 treatment. **D**. Average responses to visual stimuli of SST axons and somas. Green lines/bars: SST cIN axons, purple lines/bars: SST somas, solid lines/bars: saline treatment, dashed lines/bars: CGP-35348 treatment. (***p<0.001; mixed effects model; axons saline: mean = 6.9141, 95% CI = [6.519 7.302], axons CGP: mean = 4.99, 95% CI = [4.741 5.236], somas saline: mean = 2.39, 95% CI = [2.016 2.793], somas CGP: mean = 2.3650, 95% CI = [1.868 2.889]; error bars represent 95% confidence intervals). **E**. Example traces showing SST axonal visual response with and without NDNF cIN activation highlighting the analysis epochs. **F**. Percentage change of normalized responses of SST axons during the NDNF cIN activation epoch (0.5 sec, left; ***p<0.001, mixed effects model, saline: mean = -1.73, 95% CI = [-1.921 -1.541], CGP: mean = -0.55 95% CI = [-0.693 -0.415]) and post-activation epoch illumination (1-2 sec, right; ***p<0.001, mixed effects model, saline: mean = -36.03 95% CI = [-40.39 -31.66], CGP: mean = -24.21 95% CI = [-29.04 -19.38]). ***p<0.001, mixed effect model. Error bars represent 95% confidence intervals. **G**. Same as in **F** but for SST somas (Activation epoch: saline: mean = -0.27, 95% CI = [-0.513 -0.040], CGP: mean = -0.20, 95% CI = [-0.444 0.066]; post-activation epoch: saline: mean = -1.18, 95% CI = [-1.696 -0.688], CGP: mean = -1.10, 95% CI = [-1.679 -0.577]).

**Extended Data Fig. 8:**
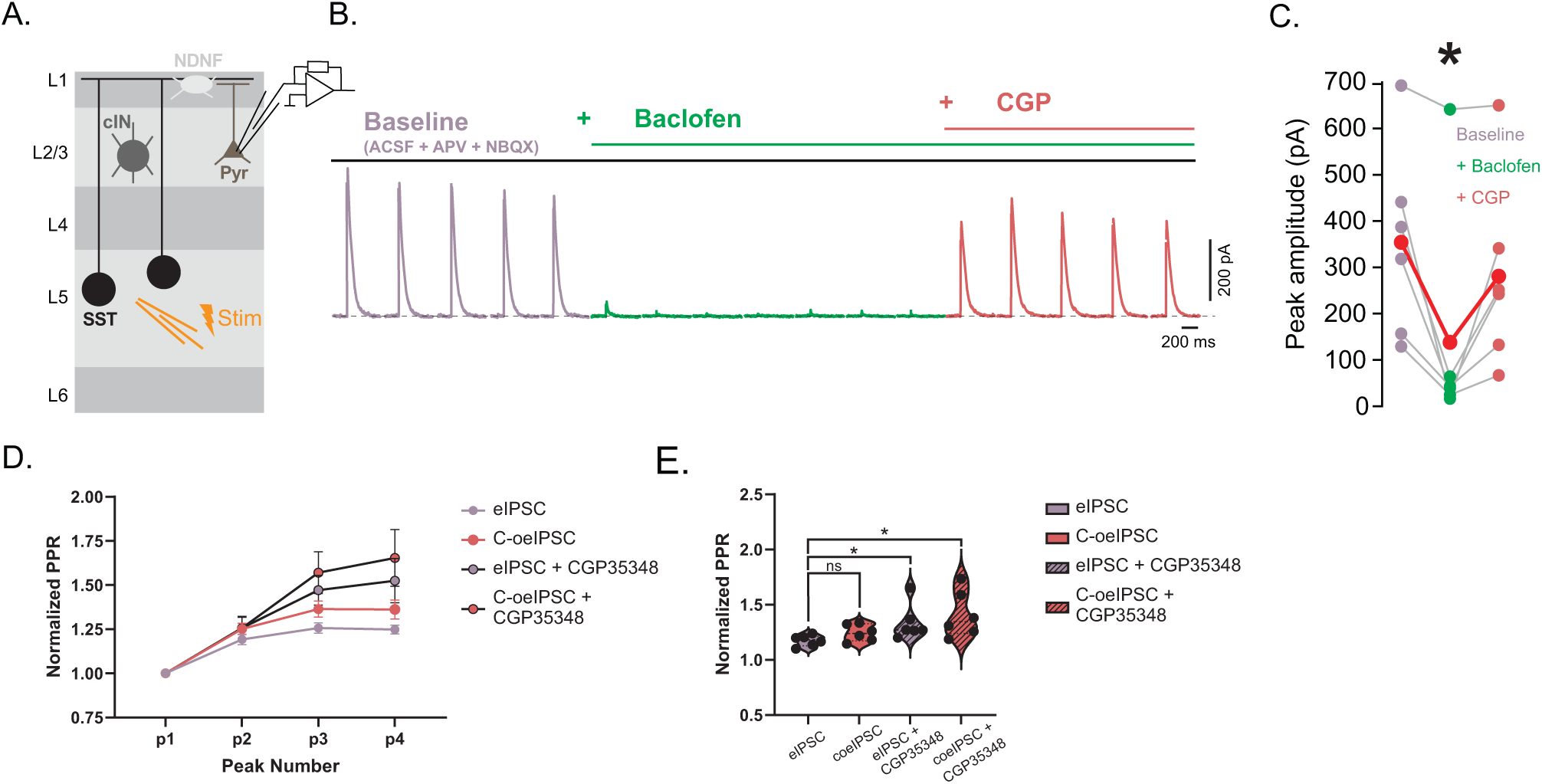
SST cIN suppression is mediated by GABAB receptors. **A.** Whole cell voltage clamp recordings of L2/3 pyramidal neurons at 0mV with translaminar electrical stimulation of L5/6. **B.** Electrical stimulation is performed in baseline ACSF (eIPSC) in the presence of excitatory blockers (D-AP5 50μM; NBQX 2μM). Electrical stimulation is repeated in the presence of Baclofen (eIPSC-Baclofen; 10μM), and Baclofen together with CGP35348 (eIPSC-Baclofen-CGP; 0.1mM) at 0.1Hz (see methods). **C.** Plot of the mean peak amplitude for the eIPSC (in purple), eIPSC-Baclofen (in green), and eIPSC-Baclofen-CGP (in peach) of each recoded L2/3 pyramidal neurons showing a significant reduction in the peak amplitude upon bath application of Baclofen (*p=0.0156 (61% reduction), n=6, Paired Wilcoxon Signed-Ranked Test), and recovery upon the addition of CGP35348 (*p=0.0156 (20% reduction), n=6, Paired Wilcoxon Signed-Ranked Test). **D.** Connected line graph showing the normalized PPR across all peaks for all cells in each condition. Error bars represent the SEM (n=6). **E.** Violin plot showing a significant increase in synaptic facilitation across all peaks between baseline electrical stimulation (eIPSC) and CGP35348 conditions (eIPSC-CGP; C-oeIPSC-CGP), as measured by the PPR (Paired Sample t-test, eIPSC vs C-oeIPSC p=0.09542; Mann-Whitney U test, eIPSC vs eIPSC-CGP *p=0.01307; Paired Sample t-test, eIPSC vs C-oeIPSC-CGP p=0.03615)

**Extended Data Fig. 9:**
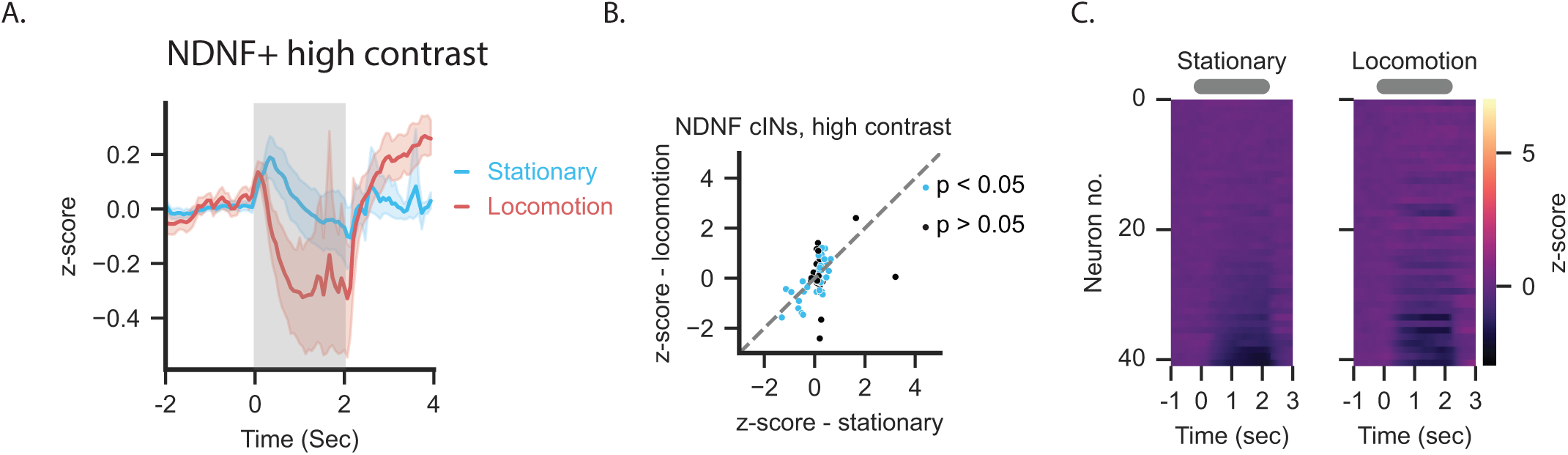
NDNF cIN responses to high contrast visual stimuli. **A**. NDNF example neuronal response to high contrast visual stimuli in stationary trials (light blue) and locomotion trials (red). **B**. Responses of individual NDNF cIN to high contrast visual stimuli while mice were stationary and to the same stimuli with locomotion at the neuron preferred direction. **C**. Normalized responses to high contrast visual stimuli of all ROIs of NDNF cINs when mice were stationary (left) and with locomotion (right; N = 2 mice, n = 89 ROIs). (n = 41, N = 2 mice). Grey shaded area and grey bar: visual stimuli. Error bars represent ±SEM.

**Extended Data Fig. 10.**
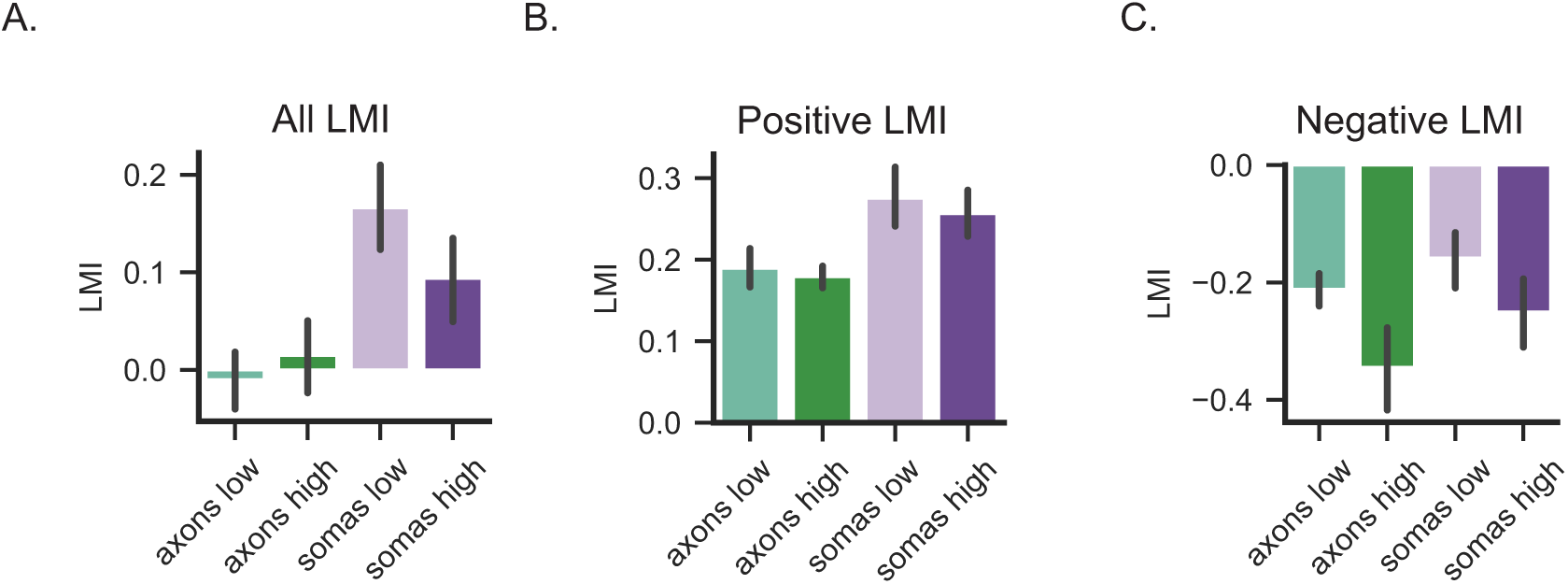
SST axons and somas responses to low and high visual stimuli in response to locomotion. **A**. Average LMI for axons presented with low contrast (light green; mean = -0.01, 95% CI [-0.04, 0.02]), axons presented with high contrast (dark green; : mean = 0.01, 95% CI [-0.02, 0.05]), somas presented with low contrast (light purple; mean = 0.16, 95% CI [0.12, 0.21]), somas presented with high contrast (dark purple; mean = 0.09, 95% CI [0.05, 0.137]). **B,** and **C**: Same as A but for positive and negative LMI respectively.

## METHODS

### Animals

Mice were bred and housed at King Abdullah University of Science and Technology (KAUST), in accordance with the guidelines of the local Institutional Animal Care and Use Committee (IACUC) protocol number LI23IACUC003, and at Cold Spring Harbor Laboratory (#2023-1264). Experiments were conducted using adult mice (older than 8 weeks) of both sexes. NDNF-dgCre or NDNF-Cre::SST-flp double-transgenic mice were generated by crossing NDNF-dgCre (Jackson laboratory, JAX No. 028536 or NDNF-Cre Jax No. 030757) mice with SST-FlpO (Jackson Laboratory, JAX No. 031629). Mice were housed in a standard 12hrs light-dark cycle and allowed ad libitum access to food and water.

### Method details Surgeries

NDNF-dgCre::SST-FlpO mice were anesthetized with isoflurane (5% for induction, and 1% maintenance during surgery) and mounted on a stereotaxic frame (David Kopf Instruments, CA). To relieve brain edema, dexamethasone (2 mg/kg) was injected Intraperitoneally thirty minutes prior to cranial window surgery. Eye cream was applied to protect the eye (GenTeal, Alcon laboratories inc, Tx). After disinfecting the scalp, lidocaine (6mg/kg) and bupivacaine (3 mg/kg) were injected under the skin and then the skin was removed. The bone surface was then cleaned with phosphate buffered saline (PBS) and the connective tissue was removed by applying hydrogen peroxide (H_2_O_2_) on the skull surface. For in vivo 2P imaging: A circular 3 mm diameter biopsy was applied to remove the skull above the visual cortex in the left hemisphere (edges of biopsy are 1 mm from the midline and 0.5 mm anterior of lambda). AAV2/9-hDlx-DIO-ChrMine-mScarlet (modified from addgene no. 130998, gift from Karl Deisseroth; titer = 4.67×10^10^ viral particles/μl) and AAV2/9-hSyn-fDIO-GCaMP8m (modified from addgene no. 162375, gift from GENIE project; titer = 3.80×10^9^ viral particles/μl) were co-injected using glass pipette with ∼30 μm diameter tip mounted using Nanoject III (Drummond Instruments Ltd.) at a speed of 1 nL/s with a total volume of 200 nL per site (2 sites; 1 mm apart diagonally). For slice electrophysiology: NDNF-dgCre::SST-FlpO or NDNF-Cre::SST-FlpO mice were injected bi-laterally with AAV-Dlx-DIO-ChrMine-mScarlet (titer: 3.50 ×10^10^ viral particles/μl) or with AAV-EF1a-DIO-Chr2-eYFP^76^; titer: >1×10^10^ viral particles/μl; addgene no. 20298) ; together with AAV-dlx-fDIO-GCaMP8m into V1 (AP:-3.5mm, ML:-/+2.5mm, DV:-0.3). After each injection, pipettes were left *in situ* for an additional 10 min to prevent backflow. In the case of *in vivo* imaging, the craniotomy was then sealed with a 3 mm glass coverslip that was glued to another 5 mm glass coverslip (Warner instruments) which in turn was glued to the skull with vetbond (Fisher scientific UK). A custom-built head-post was implanted over the exposed skull with ductile glue and cemented with C&B metabond (Parkell). Mice were monitored for three days after surgery and were injected with analgesia (Carprofen; 0.1 mg/kg of body weight).

### Two-photon Imaging

Calcium imaging data were acquired using a Brucker two-photon microscope equipped with 16x Nikon objective and an 8KHz resonant scanner. Imaging was performed using 920 nm light delivered by a Coherent Chameleon Discovery NX TPC laser system. Images were collected at a frame rate of 30 Hz for somatic imaging (2x digital zoom) and 15 Hz for axonal imaging (4x digital zoom, averaged over 2 frames).

### Visual Stimulation and Experimental Design

We generated drifting sinusoidal gratings using PsychoPy ^73^, with spatial frequency of 0.04 cycles per degree and temporal frequency of 2 Hz, at eight directions (45° steps, 16 to 40 trials per direction in a random sequence) at 100% illumination, i.e. high contrast or 20% illumination, i.e. low contrast. Stimuli had a duration of two seconds and with an inter-trial interval of four seconds. PyContol ^74^ was used to program the timing of the stimuli with precise temporal resolution and to communicate the start of the session with the imaging system via a TTL pulse.

### Processing of Calcium Imaging Data

Motion correction and region of interests (ROIs) detections (for each axonal segment and individual soma) were performed using Suite2p ^75^. Raw fluorescence traces extracted from Suite2p were further processed using custom Python scripts. To compute ΔF/F0, a baseline fluorescent signal (F0) was estimated by median filtering each trace with a 60 second window (1800 samples). ΔF/F0 was then calculated as (F-F0)/F0, where F is the original fluorescence trace. Somatic traces were then down sampled to 15 Hz to match the axonal acquisition rate, and all traces (somatic and axonic) were subsequently smoothed using a gaussian filter with a kernel width of 5 frames. Since axonal ROIs are smaller compared to somatic ROIs, we ensured that the changes in fluorescence are not due to motion artefacts in these segments. This is also evident in Fig 5 where axonal ROIs undergo opposite responses-increasing response under high contrast visual stimuli+locomotion, vs. decreasing under low contrast+locomotion.

### Optogenetic Manipulation

To simultaneously activate NDNF cINs optogenetically and record the activity of SST cINs, the red-shifted opsin ChrMine was expressed in NDNF cINs, and GCaMP8m was expressed in SST cINs. Mice were presented with high-contrast (100%) drifting gratings (same parameters as above), and in 50% of the trials, optogenetic stimulation was delivered via a 617 nm LED (1 mW) through the microscope objective at the brain surface. Using an electrotunable lens (ETL), we either imaged SST axons in L1 or SST somas in L2/3. The LED was activated for 0.5 seconds, delivered 0.5 seconds after the onset of the visual stimulus to allow for the activation of SST INs. In separate sessions, we targeted the same XY location but different depths, to record SST axons in layer 1 and SST somas in layers 2/3. Control experiments under identical conditions were performed with both ChrMine and GCaMP8m co-expressed in NDNF INs to confirm effective activation by ChrMine. In both experiments, we have kept the shutter for the photomultiplier tube (PMT) open, to allow concurrent calcium imaging and optogenetic stimulation.

### Pharmacological Manipulation

Mice received a single intraperitoneal injection of the GABA_B_ receptor antagonist CGP-35348 (0.5 mg/kg in saline, 10 mL/kg) (Tocris Bioscience, 1245/10) in order to block GABA_B_ receptor signaling pharmacologically ^61^. Each mouse was first tested following a saline injection (control), undergoing the same optogenetic stimulation protocol as described above. One hour later, allowing sufficient time for systemic drug action, the experiment was repeated following CGP-35348 administration to assess the impact of GABA_B_ receptor blockade on SST cINs responses by NDNF IN activation.

### *In vivo* Imaging Data Analysis

#### Visual stimulation analysis

To better isolate visually evoked responses from spontaneous activity, fluorescence traces of each trial were z-scored using the 2 seconds pre-stimulus period as the baseline. Subsequently, a paired t-test was performed to assess visual modulation to each presented direction by comparing the mean response during the stimulus period (2 seconds) to the mean response during the pre-stimulus baseline in stationary trials for each ROI. ROIs were considered visually modulated if they showed a significant response (p < 0.05) to at least one stimulus direction. Only modulated ROIs were included in all subsequent analyses. For population-level analysis, direction responses were normalized by aligning each ROI’s responses to its preferred direction, defined as the direction eliciting the maximum mean response during visual stimulus presentation.

#### CGP effect analysis

To normalize the CGP-35348 effect on optogenetic modulation, a percentage change was calculated for treatments with saline as well as CGP35348:

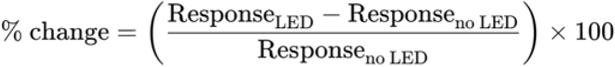

#### Locomotion analysis

Mice were head-fixed and ran freely on a belt treadmill (Labmaker) with a built-in encoder for speed and direction. The encoder information was then converted to analog voltage and was sent to the PyControl Breakout Board (Labmaker). The speed data was then converted to cm/s and smoothed post hoc. Visual stimuli trials with running speed greater than 1 cm/s and a duration longer than 2 bins (∼133 ms) were considered locomotion trials. The Locomotion modulation index (LMI) was computed to assess the responses of the neurons to locomotion which is defined as the difference between the mean ΔF/F0 during locomotion (L) trials and stationary (S) trials, normalized by the sum of both values: LMI = (L-S)/(L+S).

### In vitro Electrophysiology Experiment

*In-vitro* patch-clamp recordings were performed at least two weeks post-operation to allow viral expression in targeted cell types. Adult mice were sacrificed and the brain was removed and rapidly immersed into ice-cold, oxygenated slicing solution containing (in mM): 210 sucrose, 2.8 KCl, 2 MgSO_4_, 1.25 Na_2_HPO_4_, 26 NaHCO_3_, 6 MgCl_2_, 1 CaCl_2_, and 10 D-glucose, or an oxygenated slicing solution containing (in mM): NaCl 87, KCl 2.5, NaH2P04.H20 1.25, NaHCO3 26, Sucrose 75, D-Glucose 10, CaCl2 1, MgCl2 2, ph adjusted to 7.4 with NaOH 300µm section thick slices of the visual cortex were cut using a Leica VT1000S vibratome. Upon sectioning, the acute slices were placed into a warm, oxygenated holding solution at 32-34°C for 30 minutes, after which they were stabilized for one hour and continuously oxygenated in room temperature holding solution with the following composition (in mM): 125 NaCl, 2.5 KCl, 1.2 Na_2_HPO_4_, 25 NaHCO_3_, 2 MgCl_2_, 1 CaCl_2_ and 10 D-(+)-glucose; pH 7.4. Acute slices were then placed into the recording chamber of a SliceScope pro 1000 (Scientifica) and continuously perfused, at a rate of 4ml/min, with oxygenated artificial cerebrospinal fluid (aCSF) (in mM): 125 NaCl, 2.5 KCl, 1.25 Na_2_HPO_4_, 25 NaHCO_3_, 2 CaCl_2_, 2 MgCl_2_, and 10 glucose; pH 7.4. Slices were visualized with a camera system mounted onto an upright Olympus microscope (Orca-Flash, Hamamatsu; SciCam Pro, Scientifica; Olympus, U-TV1XC) with IR-DIC optics and a x20-x60 water immersion objective lens (XLUMPLFLN20X, LUMPlanFL, Olympus). Excitatory synaptic blockers were added to the circulating aCSF to isolate GABAergic currents during voltage clamp experiments (APV 50μM, NBQX μM; Tocris Bioscience). For experiments involving the addition of Baclofen (Tocris Bioscience), a concentration of 10μM was applied to the bath.

Patch electrodes (5-8 MΩ) were pulled from filamented thick-wall borosilicate glass (GC150F-10, Harvard Apparatus; B150-86-7.5, Sutter instrument) with a pipette puller (P-30, P-1000, Sutter instrument) and filled with cesium-based internal recording solution (in mM): Cs-methanesulfonate 130; CsCl 5; HEPES 10; EGTA.CsOH 0.2; Phosphocreatine-Tris2 8; Mg-ATP 4; Na2-GTP 0.3; QX314-Cl 5; Biocytin 0.4%.

### Electrical & Optogenetic Stimulation

Extracellular stimulation of layer 2/3 was performed using a concentric bipolar electrode attached to an electrical stimulus device (30200, FHC Inc.; ISO-Flex, A.M.P.I) or using a stimulator generator (model 2100, A-M systems) whose positive electrode was placed in the recording chamber and the negative one was inserted in a 1-2 MΩ borosilicate glass pipette positioned in L2/3 and filled with an HEPES-based aCSF containing (in mM): NaCl 125, KCl 2.5, CaCl2 2, NaH2PO4.H2O 1.25, HEPES 10, MgCl2 1, pH adjusted to 7.4 with NaOH. Concurrently, whole-cell patch clamp electrophysiology was performed on L2/3 pyramidal neurons to record the effect of electrical stimulation. All recordings were performed in the presence of NBQX (5µM) and D-AP5 (50µM) to isolate GABAergic inputs onto pyramidal neurons. Depending on the rhodopsin excitation wavelength, a green (550nm; CoolLED, pE-300white) or blue beam of light (470 nm; CoolLED, PE-4000) was sent through the objective and centered on L1 to optogenetically activate ChRmine or ChR2 expressing NDNF neurons. Recordings were performed using a Multiclamp 700B amplifier (Molecular devices) and digitized using Power1401 CED digitizer and Signal v8 software (Cambridge Electronic Devices) or an EPC10 amplifier with the software patchmaster version 2.0 (HEKA). Voltage traces were either filtered at 1kHz and sampled at 50kHz or filtered at 3kHz and sampled at 20kHz. Electrical and optogenetic activation were controlled through TTL-triggered digital output channels.

### Experimental design - Single Stimulation

The electrical stimulation was positioned on the surface of L2/3, ≥100µm away from the pyramidal cell recorded in whole cell configuration and held at 0mV. To measure the effect of NDNF cINs activation on the inhibitory inputs to pyramidal neurons, we applied 3 protocols: (1) a minimal 20 ms optogenetic stimulation (oIPSC) to evoke a robust slow-rising slow-decaying GABAergic current in pyramidal cells, (2) a minimal 1 ms electrical stimulation to evoke a robust GABAergic IPSC (eIPSC) in pyramidal cells and (3) the optogenetic stimulation immediately followed by the electrical stimulation (oeIPSC). All three protocols had 5-6 sweeps at 0.1 Hz and were repeated in a random order to reach 15-20 trials per protocol.

### Experimental design - Baclofen Experiments

To determine the involvement of GABAB receptor in the regulation of distal inhibition to L2/3 excitatory neurons, IPSCs were evoked by electrical stimulation in layer 5/6 (eIPSC). eIPSCs were pharmacologically isolated by constant bath application of APV (50 μM) and NBQX (2 μM) and L2/3 neurons were held at 0 mV. IPSCs were evoked every 10 seconds and a baseline was recorded for 5 minutes followed by a bath application of Baclofen (10 μM) for 7 minutes and a co-application of Baclofen with CGP55845 (100 μM) for 5 minutes. IPSCs from each pharmacological conditions (baseline: NBQX/APV; Baclofen: NBQX/APV/Baclofen; CGP+Baclofen: NBQX/APV/Baclofen/CGP55845) were averaged and their peak amplitude measured. To illustrate the evolution of the IPSCs across these conditions, averaged IPSCs were calculated every minute.

### Experimental design - eIPSC Train @ 20Hz

To assess the presynaptic nature of NDNF suppression of SST axonal output to pyramidal neurons, we applied three protocols under different physiological conditions. (1) a 200ms optogenetic stimulation to evoke a robust slow-rising slow-decaying GABAergic current in pyramidal neurons (oIPSC), (2) four pulses (1ms pulse width) of electrical stimulation at 20Hz to evoke a train of GABAergic IPSCs (eIPSC), and (3) optogenetic stimulation in conjunction with 20Hz electrical stimulation (oeIPSC). The 200ms optogenetic stimulation was selected to cover the entire duration of the 20Hz electrical stimulation train. All three protocols were applied at 0.06Hz for a minimum of 15-20 trials per protocol. This sequence was applied in three different recording ACSF conditions. (I) A recording ACSF with 2mM Ca^2+^, (II) A low Ca^2+^ recording ACSF with 1mM Ca^2+^, (III) A low Ca^2+^ recording ACSF and 100μM CGP-35348 (Tocris Bioscience). A minimum waiting period of three minutes was observed after changing ACSF conditions to allow the neuron to adapt to new physiological conditions before recording.

### Electrophysiology Data Analysis

To compare IPSC peak amplitude evoked by the L2/3 electrical stimulation with (oeIPSC) and without the optogenetic activation of NDNF neurons (eIPSC), we normalized each trace to its baseline and subtracted the NDNF component (oIPSC) from the oeIPSCs. For all pyramidal cells, the cumulative sum of each individual oIPSC trace is measured during the 20 ms of the optogenetic stimulation. This time window contains the initial rising phase of the NDNF component, and measuring its integral depicts its kinetic. The same operation is done on each oeIPSC, and the oIPSC with the closest rising phase kinetic is subtracted from it. Corrected oeIPSCs (C-oeIPSCs) and eIPSCs are then averaged, and their peak amplitude measured.

### Normalized PPR Analysis and Selection Criteria

The normalized paired-pulse ratio (PPR) for each cell was calculated by averaging the trials for each condition, and dividing each peak of the 20Hz electrical stimulation train by the amplitude of the first peak of the condition. If the average ratio across all four peaks is greater than one, the synapse is considered to be facilitating. If the average ratio across all four peaks is less than one, the synapse is considered to be depressing.

### Rate of Change analysis

The normalized PPR for each condition was plotted, a tangent line was fitted, and slope calculated at peak2 to evaluate the rate of change between peak1 and peak2 (Tangent v1.82, OriginLab 2022). The greater the normalized PPR change between peak1 and peak2, the greater the slope value.

### Peak amplitude subtraction

Current traces for each condition were initially normalized to their baseline and grouped by recording condition. Next, the peak amplitude values of the corrected traces from the simultaneous optogenetic & electrical stimulation (C-oeIPSC) were subtracted from the peak amplitude values of the electrical stimulation (eIPSC) train in a pairwise fashion.

### Perfusion and tissue preparations

For all histological experiments, mice were deeply anesthetized via intraperitoneal injection of lethabarb (60 mg/ml, intraperitoneally) and transcardially perfused with 1× phosphate-buffered saline (PBS), followed by 4% paraformaldehyde (PFA) in 1× PBS. Brains were carefully extracted and post-fixed overnight at 4°C, for 2 hours at room temperature or for 4 hours at 4°C. Following fixation, tissues were either cryoprotected by immersion in 30% sucrose in 1× PBS until fully equilibrated and sectioned on freezing sliding microtome (Leica, SM2010R) or sectioned on a vibratome (Leica VT1000S), without cryoprotection

### Immunohistochemistry

Free-floating brain sections were incubated in blocking solution (10% normal donkey serum, 0.3% Triton X-100 in 1X PBS) at room temperature for 1 hour, followed by incubation in primary antibody diluted in blocking solution at 4°C overnight. The following day, sections were rinsed in 1X PBS for 10 minutes 3 times, followed by secondary antibody incubation in the same blocking solution for 1 hour at room temperature. Sections were then rinsed in 1X PBS for 10 minutes 3 times. Sections were counterstained with DAPI (5 μM in 1x PBS,) for 3 minutes and mounted with fluoromount-G (Invitrogen, 00-4958-02). Primary antibodies used in this study include Chicken-anti-GFP (Abcam, ab13970, 1:2000), Guinea pig-anti-RFP (Synaptic Systems, 390 004, 1:1000), Anti-GABABR1 (NeuroMab, N93A/49, 1:500), and SV2C (SYSY, 119 202,1:200). Secondary antibodies (diluted at 1:500) used in this study include Alexa Fluor 488 Goat anti-Chicken (Invitrogen, A11039) and Alexa Fluor 594 Goat anti-Guinea Pig (Invitrogen, A11076).

### Super-resolution confocal imaging

Fluorescent images were acquired on a Zeiss LSM 980 confocal microscope equipped with Airyscan2 detection. Samples were imaged using a 63× oil-immersion objective. Excitation was achieved with 405, 488, 555, and 647 nm laser lines to detect the respective fluorophores. Z-stacks were collected with optimal step sizes determined by the ZEN software to cover full depth. Airyscan processing was applied using the manufacturer’s recommended settings in ZEN software to improve resolution and signal-to-noise ratio. Identical imaging parameters were maintained across experimental groups, and all post-processing was limited to Airyscan reconstruction.

### Histological image analysis and quantification

Synapses and 3D reconstructions using Imaris v10.2 (Oxford Instruments) were quantified from confocal z-stack images acquired using AiryScan. The confocal images were subjected to 3D reconstruction to visualize and analyze multiple neuronal components, including SST cINs, NDNF cINs, GABAB1 receptors, and SV2C using Imaris 10.2.0 software. Images were processed with a Gaussian filter (width: 0.0353 μm) and background subtraction (11 μm).

Surface-based analysis was performed as follows: Neuronal surfaces were created for each channel to demarcate SST, GABAB1R, NDNF, and SV2C distributions using absolute intensity thresholds set to accurately identify neurons of interest. Channel masks were generated from these surfaces to identify co-occurrence regions of GABAB1R+ SST and SV2C+ NDNF axons. Quantitative colocalization analysis was performed using Imaris’s “Colocalization” module to assess GABAB1 receptor distribution in SST neurons and SV2C distribution in NDNF neurons. Summary and detailed statistics were collected for the new GABAB1R+ SST and SV2C+ NDNF channels, and areas were determined by surface.

Distance measurements were calculated from each GABAB1R+ SST spot object created from the previous mask surfaces of GABAB1R+ SSTand SV2C+ NDNF axons. Spatial proximity cumulative distribution analysis was performed at 1 μm, 2 μm, and 3 μm intervals. The quantification was performed by applying distance-based filters to categorize spatial relationships into three proximity classes: proximity (<1 μm), intermediate distance (1-2 μm), and distant (>2 μm).

Individual views of the distribution of each object from GABAB1R+ SST and SV2C+ NDNF were acquired with X, Y, and Z coordinates using the Vantage Wizard in IMARIS, enabling quantitative assessment of spatial clustering and potential synaptic proximity between GABAB1R+ SST and SV2C+ NDNF populations for unbiased analysis. Euclidean distances between each GABAB1R+ SST (obj1) and every individual SV2C+ NDNF (obj2) object per stack were calculated as follow:

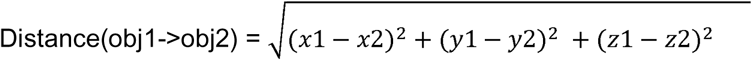

With GABAB1R+ SST and SV2C+ NDNF objects having euclidean coordinates (x1,y1,z1). To evaluate an unbiased density of contact between the two objects, these distances were binned at 1 micron from 0-10 microns and the number of events per bin, per GABAB1R+ SST object has been calculated. The number of SV2C+ NDNF objects increases linearly with the volume of a sphere centered on GABAB1R+ SST objects, thus, each bin has to be normalized by the volume of their corresponding hollow-sphere to account for this increased probability. The volume of each hollow-sphere corresponding to each bins of 1 micron as been calculated as follow: Volume hollow-sphere = ((4÷3) × *π* × *rb*^3^) − ((4÷3)) × *π* × *ra*^3^) with “ra” and “rb” the radius of the respective sphere for bins n-1 and n. For the first bin (0-1 micron), a sphere volume was used. Finally, the density of GABAB1R+ SST > SV2C+ NDNF objects per bin has been averaged. The averaged voxel size being 0.065 × 0.065 × 0.144 micron, we re-ran the same analysis between 0-2 microns with a binning of 0.2 micron to have a better resolution of the GABAB1R+ SST -> SV2C+ NDNF objects density within the canonical synaptic space.

To validate the co-occurrence rate between markers, apposition analysis was performed between SST and GABAB1R distributions. Cluster centers and areas were retrieved in original and randomized stacks (GABAB1R vertically flipped), which yielded close to 0% occurrence, confirming the specificity of the original analysis.

### Single Molecule Fluorescent In Situ Hybridization (smFISH): RNAscope

Brains were cryoprotected in 20 % followed by 30 % sucrose, embedded in OCT and stored at −80 °C; coronal cryosections (30 µm) were cut at −20 °C, mounted on SuperFrost Plus slides (Fisher Scientific, 12-550-15), air-dried (1 h, −20 °C) and baked (30 min, 60 °C). Sections were post-fixed in 4 % PFA (15 min), rinsed in PBS, dehydrated through graded ethanols (50 %, 70 %, 2 × 100 %; 5 min each) and air-dried (5 min); a hydrophobic barrier was drawn with an ImmEdge pen (Vector, H-4000) and dried (5 min). Endogenous peroxidase activity was quenched with RNAscope Hydrogen Peroxide (ACD, 322381) for 10 min at room temperature (RT), followed by target retrieval in pre-boiled 1 × Target Retrieval Reagent (ACD, 322000; 5 min, ≥99 °C), a 3-min 100 % ethanol dip and drying (5 min, 60°C). Protease permeabilization was performed with Protease III (ACD, 322381; 30 min, 40°C in a HybEZ II oven, 321711), then slides were rinsed twice in distilled water. Probe-Mm-Ndnf-C1 (ACD, 447471) and Probe-Mm-Sst-C3 (ACD, 404631-C3) were hybridized for 2 h at 40 °C; sections were washed twice in 1 × Wash Buffer (ACD, 310091) and incubated sequentially with AMP-1, AMP-2 and AMP-3 (30 min at 40 °C, respectively) with identical washes between steps. Signal was developed in three cycles: HRP-C1 (15 min, 40 °C) followed by Opal 520 (Akoya Biosciences, FP1488001KT; 1:1 000 in TSA buffer, 20 min, 40°C) and HRP blocker (20 min, 40 °C), then analogous HRP-C3 with Opal 620 (Akoya Biosciences, FP1495001KT), each separated by washes in 1 × Wash Buffer. Nuclei were counterstained with RNAscope DAPI (included in 323110; 30 s, RT), and sections mounted in ProLong Gold Antifade Mountant (Thermo Fisher, P36930) and cured overnight in the dark. *Confocal Imaging:* Images were acquired using a Leica Sp8 confocal microscope with either a 40× or 63x objective.

### Single Molecule Fluorescent In Situ Hybridization (smFISH): HCR

Tissue samples were prepared for in situ hybridization. Samples were pre-incubated in HCR™ HiFi Probe Hybridization Buffer at 37 °C for 30 min, followed by incubation with an HCR™ v3 probe set targeting Lamp5 in hybridization buffer at 37 °C for >3 h. Samples were washed 4 × 15 min with pre-warmed HCR™ HiFi Probe Wash Buffer (1×, diluted from 4× stock).

Signal amplification was performed using HCR™ Gold reagents. Samples were pre-incubated in Amplifier Buffer at room temperature for 30 min. Hairpins (h1 and h2) were snap-cooled by heating to 95 °C for 90 s and cooling to room temperature in the dark for 30 min, then added to Amplifier Buffer to prepare the amplification solution. Samples were incubated in amplification solution at room temperature in the dark for >3 h, followed by 4 × 15 min washes in 1× HCR™ Gold Amplifier Wash Buffer. After amplification, samples were stored at 4 °C protected from light until imaging and mounted in antifade medium with or without DAPI counterstaining. Images were acquired with Zeiss LSM 780 with either a 40× or 63x objective.

### Quantification of viral expression across cortical layers

Layer boundaries in the visual cortex were determined by visually assessing cell densities across layers. Subsequently, neurons expressing the reporter gene were manually quantified in ImageJ software and then normalized to DAPI stained cells.

Specificity of the viral expression was assisted by counting the cell expressing the reported gene and cells co-labeled with the respective RNAscope marker following the formula:

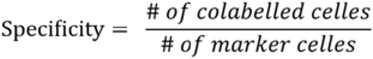

### Electron microscopy (EM) analysis

Analysis of axo-axonal connectivity was performed on the serial section transmission EM (TEM) volume of mouse visual cortex acquired as part of the MICrONS project (Bae et al., 2025; Schneider-Mizell et al., 2025). Using the Connectome Annotation Versioning Engine CAVEclient (https://tutorial.microns-explorer.org/quickstart_notebooks/01-caveclient-setup.htm) in Python v3.10.18 with the latest released version of the public dataset (v1412), which contained cell-type and connectivity annotations from (Schneider-Mizell et al., 2025) based on inhibitory neuron circuit connectivity from a connectomic census. We queried Distal Targeting Cells (DistTC), Sparsely Targeting Cells (SparTC) and Inhibitory Targeting Cells (InhTC), which were associated with Martinotti or Somatostatin cells, Neurogliaform/NDNF cells and Bipolar/ VIP disinhibitory neurons, respectively. For Layer 1 focus, we selected SparTC and InhTC based on their soma depth. L1 was set as 0 and soma depth measured in um (soma depth (um) = Y(distance in voxels) × 0.004 (scale um) - 300 um offset), as defined from plotting the Y distance distribution of all cell types, including L4 excitatory cells (‘4P’). SparTC and InhTC cells were filtered to lie within the top 250 um from the surface and selected a total of 547 and 733 respectively. In the same study, synapse detection and cell assignment were performed with a trained convolutional network. The number of conventional synapses made by SparTC and InhTC onto DistTC was extracted. This number served as the baseline detection of conventional synapses onto Martinotti cells. However, prediction network methods account for conventional synapse orientation as a marker of synaptic cleft detection (Parag et al., 2019; Turner et al., 2020), therefore precluding the detection of axo-axonic interactions. These atypical interactions were manually identified with contact and presence of presynaptic vesicles in InhTC or SparTC. The previous annotations may not have been fully validated, as we observed that some annotated DisTC contained misclassified cells, including pyramidal excitatory neurons. We therefore picked single DisTC based on their morphology. 3D Mesh reconstruction was always in full, especially axonal arborization. We therefore preferred DisTC with full extension of axons within L1 and of typical Fanning out- and T-shapes. “Pt_root_id” unique IDs of identified L1-SparTC or L1-InhTC were displayed in the modified visualization tool, Neuroglancer (ngl.microns-explorer.org), with the public segmentation dataset (minnie65) and corresponding precomputed mesh. SparTC and InhTC identity was blinded to the investigator. Final annotations of synapses and contact for all DistTC included in the study are in Supplementary Table 1 within corresponding Neuroglancer links. An example of one synapse for each cell is presented into a video (Supplementary videos #1-8).

### Statistical Analysis

Data are represented as mean ± 95% confidence intervals (*in vivo*) or mean ± SEM for the rest. All statistical analyses were computed using Python (version 3.12.6 and 3.10.18), using Panda, Numpy and MatPlotLib libraries. Plots were generated with custom Python scripts, Prism 8 and 10 (GraphPad) and OriginPro (OriginLab 2022). See statistics table for details of tests used and exact p values.

### Data and code availability

Statistical analysis of all data is available in Supplementary Table 2. Code to analyze *in vivo* data is available on GitHub.

